# Species tree branch length estimation despite incomplete lineage sorting, duplication, and loss

**DOI:** 10.1101/2025.02.20.639320

**Authors:** Yasamin Tabatabaee, Chao Zhang, Shayesteh Arasti, Siavash Mirarab

## Abstract

Phylogenetic branch lengths are essential for many analyses, such as estimating divergence times, analyzing rate changes, and studying adaptation. However, true gene tree heterogeneity due to incomplete lineage sorting (ILS), gene duplication and loss (GDL), and horizontal gene transfer (HGT) can complicate the estimation of species tree branch lengths. While several tools exist for estimating the topology of a species tree addressing various causes of gene tree discordance, much less attention has been paid to branch length estimation on multi-locus datasets. For single-copy gene trees, some methods are available that summarize gene tree branch lengths onto a species tree, including coalescent-based methods that account for heterogeneity due to ILS. However, no such branch length estimation method exists for multi-copy gene family trees that have evolved with gene duplication and loss. To address this gap, we introduce the CASTLES-Pro algorithm for estimating species tree branch lengths while accounting for both GDL and ILS. CASTLES-Pro improves on the existing coalescent-based branch length estimation method CASTLES by increasing its accuracy for single-copy gene trees and extends it to handle multi-copy ones. Our simulation studies show that CASTLES-Pro is generally more accurate than alternatives, eliminating the systematic bias toward overestimating terminal branch lengths often observed when using concatenation. Moreover, while not theoretically designed for HGT, we show that CASTLES-Pro maintains relatively high accuracy under high rates of random HGT.

**Code availability:** CASTLES-Pro is implemented inside the software package ASTER, available at https://github.com/chaoszhang/ASTER.

**Data availability:** The datasets and scripts used in this study are available at https://github.com/ytabatabaee/CASTLES-Pro-paper.

## Introduction

Summarizing a collection of potentially conflicting trees inferred from different parts of the genome (i.e., gene trees) to obtain a species tree has now become a routine analysis. This approach promises to account for biological processes such as incomplete lineage sorting (ILS), gene duplication and loss (GDL), and horizontal gene transfer (HGT) that create discordance between gene trees and the species tree (Maddison, 1997). Prior studies have confirmed that the most accurate methods for estimating the topology of species trees are those that take biological sources of heterogeneity into account (Jiang et al., 2020; Kubatko and Degnan, 2007; Molloy and Warnow, 2018). Several methods use likelihood under a model of genome evolution coupled with Bayesian MCMC inference to jointly estimate both topology and branch lengths of gene trees and species trees. These methods tend to be accurate, but they are computationally intensive. One alternative is the two-step approach of inferring gene trees independently and then using a summary method to build a species tree. This alternative has been more scalable and generally accurate (Mirarab et al., 2021), spurring the development of many such methods (e.g., Larget et al., 2010; Liu and Yu, 2011; Liu et al., 2009; Solís-Lemus and Ané, 2016; Vachaspati and Warnow, 2015; Wang and Nakhleh, 2018). Some of these summary methods (e.g., Chaudhary et al., 2013; Legried et al., 2021; Molloy and Warnow, 2020; Wehe et al., 2008; Willson et al., 2022; Zhang et al., 2020) account for GDL and can take as input multi-copy gene trees, vastly expanding the set of loci that can be used (Smith and Hahn, 2021). For example, the ASTRAL family of methods (now available in the ASTER software package) use variants of the median tree problem based on the quartet distance (Mirarab et al., 2014; Zhang and Mirarab, 2022b; Zhang et al., 2018) and have been extended to multi-copy input trees (Zhang et al., 2020). The ASTRAL family is widely used, including the ASTRAL-pro extension to multi-copy input (e.g., Chanderbali et al., 2022; Ding et al., 2023; Guo et al., 2021; Li et al., 2024).

Species trees are most useful if they are furnished with branch lengths, as many downstream applications, including dating, comparative genomics, and the study of diversification and adaptation, depend on branch lengths. However, widely used summary methods such as ASTRAL do not produce the branch lengths needed for downstream analysis. Species trees can be furnished with coalescent unit (CU) lengths only for internal branches (Sayyari and Mirarab, 2016) (unless multiple individuals are available), and GDL-based methods often produce no branch length. Meanwhile, downstream applications often require branch lengths in the unit of either substitution per site (SU) or time. The standard *ad-hoc* solution is to estimate the species tree topology using a summary method and then infer the branch lengths using concatenation. Often, branch lengths are optimized on a fixed topology using maximum likelihood applied to a concatenation of all genes (e.g., Jarvis et al., 2014; Song et al., 2012; Zhu et al., 2019). An alternative is using distance-based approaches, such as ERaBLE (Binet et al., 2016) and TCMM (Arasti et al., 2024), to summarize patristic distances from gene trees onto the species tree. Neither concatenation nor distance-based methods directly model the biological processes that create gene tree heterogeneity, reducing their theoretical justification. Nevertheless, these methods have the potential advantage of being agnostic to the source of discordance. Ultimately, which approach should be preferred is an empirical question with important downstream implications (Moody et al., 2022b).

We recently introduced the CASTLES (Tabatabaee et al., 2023) method for estimating SU branch length for a fixed species tree topology, specifically designed to handle ILS, as modeled by the multi-species coalescent (MSC) model. CASTLES estimates species divergence times as opposed to *genic* divergences, which are expected to be older (Edwards and Beerli, 2000). CASTLES had higher accuracy than alternatives in our simulations. Nevertheless, it has several limitations. Most importantly, CASTLES is limited to single-copy gene trees and, therefore, cannot be used with multi-copy input trees, severely limiting its applicability. To our knowledge, the only method that can estimate SU branch lengths from multi-copy gene trees is Species-Rax (Morel et al., 2022), which does model GDL but does not model ILS and, thus, deep coalescence. In addition, the study by Willson et al. (2022) shows that SpeciesRax can be less accurate than ASTRAL-Pro and other methods in conditions with ILS and is also less scalable. Even the use of concatenation in the presence of GDL is complicated and requires additional techniques, such as DISCO (Willson et al., 2022), to decompose multi-copy genes into single-copy ones. Beyond the lack of support for GDL, CASTLES was not tested under conditions with HGT, and even for ILS, it required several approximations that could reduce its accuracy.

In this article, we dramatically advance the CASTLES methodology to address its major limitations and broaden the scope of conditions under which it is tested. We present a dynamic programming algorithm for estimating branch lengths of a species tree from multi-copy gene family trees that have evolved with GDL in addition to ILS, leading to a new method called CASTLES-Pro. In addition, we improve upon CASTLES by relaxing some approximations and modifying other assumptions. Beyond ILS and GDL, for which it is designed, we use simulations to test how CASTLES-Pro performs under conditions that include substantial levels of ILS and HGT. In simulations, we show that the method is accurate, robust to various sources of heterogeneity, and scalable to thousands of species and genes. On diverse biological data ranging from the root of the tree of life to recent speciations, we show that using CASLTES-Pro instead of concatenation dramatically alters branch lengths. We have incorporated CASTLES-Pro inside the ASTER package of tools, providing a new C++ implementation compared to CASTLES. Thus, any user of ASTRAL-Pro or ASTRAL-IV would automatically obtain SU branch length with no additional step needed.

## Material and Methods

### CASTLES-Pro Algorithm

We first review the CASTLES algorithm, on which CASTLES-Pro is based. We then describe how it is extended to handle multi-copy gene family trees and end by explaining the methods used to enhance CASTLES for both single-copy and multi-copy gene trees.

### CASTLES

The input to CASTLES is a species tree topology and a set of *k* single-copy gene trees with branch lengths in units of the expected number of substitutions per site (SU). Its output is the species tree with SU lengths on all branches. The model it assumes is parametrized by a species tree topology, and for each branch *i*, we are given the CU length (*T*_*i*_) and per-branch substitution rate (*µ*_*i*_) in the unit of substitutions per site per CU.

Thus, *µ*_*i*_ increases not only with per generation substitution rate *but also* with the effective population size, *N*_*e*_. Species branch lengths are multiplied by corresponding substitution rates to give the species tree SU length *t*_*i*_ = *T*_*i*_*µ*_*i*_. We use *t* = ⟨*t*_*a*_, *t*_*b*_, …⟩ and *µ* = ⟨*µ*_*a*_, *µ*_*b*_, …⟩ as shorthand. The CU gene trees are generated from the species tree using the MSC model. Then, each gene tree branch length is scaled by mutation rates corresponding to all species tree branches that it traverses.

CASTLES is based on expected values of gene tree branch lengths under the MSC. Treating species tree branch lengths *t* and mutation rates *µ* as unknown parameters, we can analytically calculate the expected length of the gene tree branches. For a simple cherry tree (a,b):*T* with mutation rate *µ*_*a*_, *µ*_*b*_ for terminal branches and *µ*_*r*_ for the parent branch, this expected length is

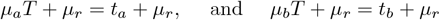

for terminal branches of *a* and *b*, resp. Note that each terminal length has an extra *µ*_*r*_ term. This gap between speciation and genic divergence (1 CU in expectation; i.e., *N*_*e*_ generations) is not modeled by concatenation or methods that do not account for coalescence and can impact downstream analyses such as dating.

We can extend this idea to quartet trees using more advanced calculations and distinguishing gene trees that match or do not match the species tree. These equations would be of the form *E*(*L*) = *f* (*t, µ*) where *L* is a random variable representing the length of a terminal (*L*_*T*_) or internal (*L*_*I*_) branch of a quartet gene tree matching the topology of the species tree or conflicting with it (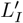 and 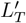). We derived those equations, reproduced in Tables S4 to S6. To estimate species tree SU lengths given a set of gene trees, we first compute the average branch lengths for gene trees that match or conflict with the topology of the species tree 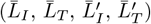; see Figure 1A. Equating the theoretical expected values (with unknown parameters) with observed values, we get a set of equations that can be analytically solved. CASTLES employs several simplifying assumptions to reduce the number of parameters further and simplify the equations (Table S5). For example, to estimate an internal branch with length *t*_1_ = *T*_1_*µ*_1_ (Fig. 1B), we obtain

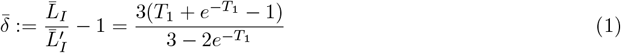

which we solve for *T*_1_ to obtain

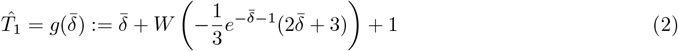

where *W* (.) is the Lambert *W* function. We then estimate substitution rates *µ*_1_ using a second equation with further simplifying assumptions (see Supplementary Section B) to obtain 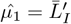 and thus:

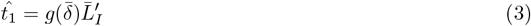

**Figure 1:**
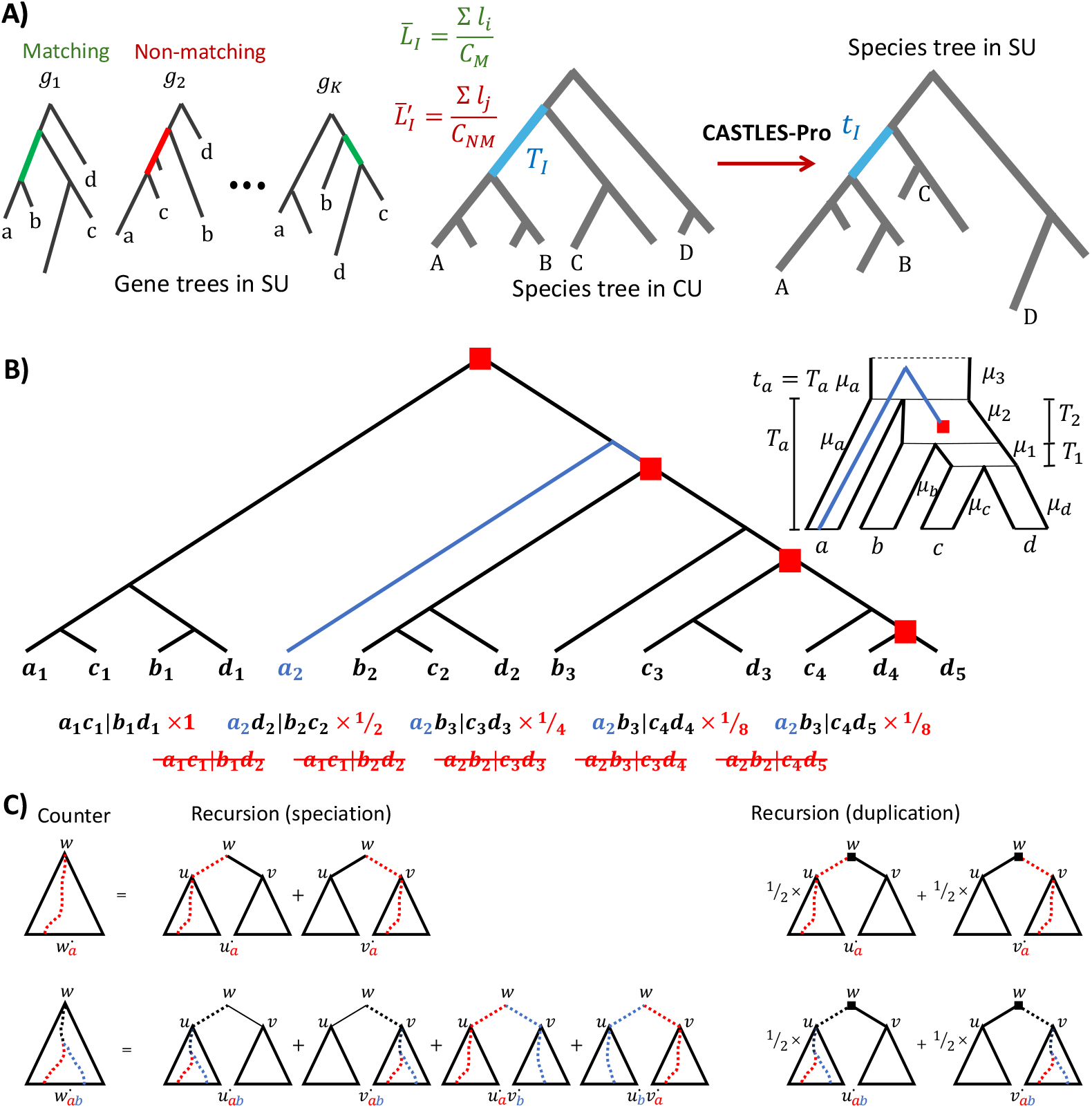
A) Illustration of matching and non-matching gene trees and the CASTLES-Pro algorithm. The SU length of each species tree branch is calculated using formulas derived from the average SU lengths of that branch in quartets induced from gene trees that either match or do not match the topology of the species tree. B) Illustration of an unbalanced species tree and a gene family tree undergoing consecutive duplication events noted in red boxes. Each species tree branch has a substitution rate *µ*_*i*_; branches of the gene tree inherit rates of species tree branches they pass through (e.g., the *a*_2_ branch will inherit *µ*_*a*_ and *µ*_2_). We show all quartet trees involving only orthologous genes alongside their respective weights, followed by examples of quartet trees that include paralogous genes (not counted). The weight of a quartet is 2^−*d*^, where *d* is the number of duplication events the quartet passes through. The sum of all quartets, including a branch (e.g., *a*_2_), is at most 1 but can be lower (e.g., ^1^*/*_2_ for *b*_3_). C) Examples of counters for computing the weighted count of gene tree quartets using dynamic programming (see Supplementary Fig. S1 for full results). For the species tree quadripartition *AB*|*CD*, we compute the mean length of the branches matching (or not matching) it in a bottom-up traversal. At each node *w*, several counters are updated; the two simplest counters are shown: 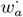 (weighted number of leaves corresponding to *a* ∈ *A* below *w*) and 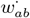 (weighted number of pairs (*a, b*) ∈ *A × B* with a speciation node as LCA at or below *w*). Note the weights for duplication.

CASTLES extends these calculations to *n >* 4 species by computing averages over all quartets around each species tree branch, a task that can be done in *O*(*n*^2^) using a dynamic programming algorithm.

### Handling Duplication and Loss

Gene duplications create quartet trees with paralogous gene copies. CASTLES-Pro addresses this issue by striving to exclusively use quartets devoid of paralogous genes. Following ASTRAL-Pro2, we first tag each internal node of each input gene tree as either a duplication or a speciation event, and we assume these *tags* are accurate (in practice, they are obtained using parsimony and may have errors). CASTLES-Pro operates similarly to CASTLES, with two major changes. The main change is that a quartet contributes to empirical mean branch length 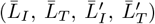 only if the least common ancestors (LCAs) of all pairs of its leaves are speciation nodes (these are called orthologous quartets). If all the tags are correct, orthologous quartets will follow the MSC expectations (Zhang et al., 2020) and, thus, CASTLES assumptions.

The second change relates to the potentially uneven rates of duplication across gene families, which can lead to certain branches being overrepresented in the final means. Figure 1B demonstrates an example; four out of five orthologous quartets share the *a*_2_ branch indicated in blue; however, the duplication events are on a separate branch and result in using *a*_2_ four times in computing mean branch lengths of *a* (e.g., 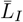) for no apparent reason. Counting all orthologous quartets without weights can increase the impact of individual branches and, thus, the variance in estimated means. We use a weighting scheme to mitigate the impact of this overrepresentation. The weights are simply set to 2^−*d*^ where *d* is the number of duplication nodes falling on the subtree spanned by the quartet tree. With this scheme, it is easy to show that the total weight of quartets that include a branch will not exceed 1 in any gene family tree.

Beyond eliminating obvious paralogs and weighting ortholog quartets, the main challenge is computing mean branch lengths efficiently instead of the trivial *O*(*n*^4^) algorithm that lists all quartets. We designed a dynamic programming algorithm that achieves this goal in *O*(*n*^2^) time (see Supplementary Section A and Algorithm S1). Compared to CASTLES, we needed to refine the set of our counters updated in the dynamic programming (demonstrated in Supplementary Figs. S1 and S2.) Crucially, at duplication nodes, the recursive formulas change to ignore non-orthologous quartets and implement the weights (see Fig. 1B). After calculating the means, CASTLES-Pro assigns lengths to each branch of the species tree in an *O*(*n*) pre-order traversal of the tree, and therefore the total runtime of the CASTLES-Pro algorithm is *O*(*n*^2^).

### Better approximations, assumptions, and handling of short branches

Even for single-copy gene trees, CASTLES-Pro improves on CASTLES in three ways and, as we will see, dominates it in terms of accuracy (thus, CASTLES-Pro replaces CASTLES.) One change is related to a simplifying assumption: CASTLES-Pro uses a slightly different approach for handling dependencies on the parent branch when calculating the length of a terminal branch of a cherry. Due to space limits, this change is explained in supplementary material (Sec. B). We elaborate here on the other two changes.

CASTLES employed a weak approximation that we eliminate in CASTLES-Pro. Instead of directly using the Lambert function *W* in Eq. (2), CASTLES used a Taylor approximation to get 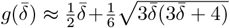. However, this approximation underestimates the true value as we move away from its focal point (0), causing a systematic underestimation bias (Fig. S4). Oddly, our simulation studies show that the Taylor approximation works better in practice on estimated gene trees despite this bias. Attributing this odd observation to difficulties with numerical precision, in CASTLES, we opted to use the Taylor expansion. However, we have now discovered a different explanation for this pattern.

Our simulations show that when using true gene trees, the Lambert *W* function is superior, whereas Taylor gradually becomes better as gene tree estimation error (GTEE) increases (Fig. S5). Taylor’s better performance is due to the interplay between two opposing sources of bias. As GTEE increases, we tend to overestimate 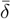, a pattern that can be explained. With a low phylogenetic signal and hence high GTEE, many of the short internal branches, which are more prevalent among gene trees not matching the species tree, become zero-event (i.e., record no substitutions). The inferred length of these supershort branches is often driven by a pseudocount used in the maximum likelihood tools (e.g., 1e-6), which is often an underestimation of the true value. Since 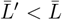 to begin with, these underestimations happen more for 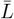 than 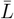, and thus, 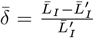 tends to get overestimated, which in turn offsets the underestimation bias of the Taylor approximation. This lucky canceling is not enjoyed if we use the exact Lambert W function. As the signal increases, or with true gene trees, there is no overestimation to offset Taylor’s bias, leading to the exact Lambert equation working better.

In CASTLES-Pro, we directly address the overestimation of 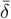 under high GTEE conditions and switch to using the exact Lambert W function in Eq. (2). If the length of an alignment is *s*, a branch of length 1*/s* would expect to see one substitution. Thus, branches substantially below 1*/s* are often zero-event and underestimated by a pseudocount. To address this, we simply add a psuedocount of 1*/s* to both 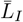 and 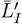, obtaining an adjusted value for *δ* given by 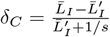. This is, in principle, similar to adding a pseudocount to binomial parameter estimation, which is equivalent to a posterior estimate under a Dirichlet prior. As sequence length increases and GTEE decreases, so does the pseudocount (in the limit, *s* = ∞ for true gene trees, giving a zero pseudocount). The value of *s* can be adjusted by the user (default: 1000).

Another difficulty arises when 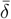 is negative, indicating that the average length in non-matching gene trees is larger than that of matching gene trees. This is unexpected under MSC and will not happen with infinitely many error-free gene trees; in practice, however, it can happen for many reasons. CASTLES simply resorted to replacing 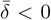 with a fixed pseudocount of *δ*_*p*_ := 10^−3^. For some causes of negative 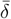, this is not a good approach. When a branch is very long with a low level of ILS, there are very few non-matching gene trees. Furthermore, these non-matching gene trees can differ from the species tree due to reasons other than ILS, such as paralogy, horizontal transfer, incorrect homology, etc. Thus, the (few) non-matching gene trees can have average lengths that are larger than the average length of matching gene trees, leading to a negative 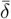. In such cases, simply using the mean of matching gene trees 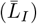 is a better approximation. In contrast, the CASTLES approach of using a small pseudocount *δ*_*p*_ in the original equation 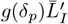 makes sense for very short branches. CASTLES-Pro handles negative 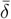 using a formula that takes the level of ILS into account and transitions between these two appraoches. For 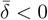 we use:

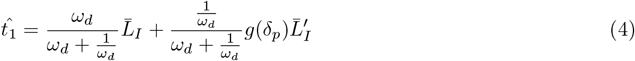

where *ω*_*d*_ = *log*_10_(*k*)*d* is the weight of the two formulas; *d* is the quartet-based CU length of the branch (see Sayyari and Mirarab, 2016) and *k* is the number of gene trees. As gene tree discordance decreases, *d* and *ω*_*d*_ increase and in the limit, 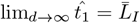 in Eq. (4); this is justified because for long branches, the deep coalescence has a *relatively* small impact compared to the full length. For short branches, *d* decreases, and in the limit 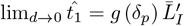, we resort to the original formula (3) used with the pseudocount. Thus, we transition from relying on average matching gene tree length 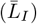 for branches with little discordance to our original estimate for high discordance; the rate of transitioning between the two approaches is governed by the number of genes, with more genes leading to faster adoption of 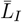; this is because the discordance-based estimates of CU length (*d*) are more accurate with more genes.

### Experimental Study

We compare CASTLES-Pro to other branch length estimation methods using three sets of simulated datasets and nine published biological datasets with gene tree discordance due to ILS, GDL, and HGT (Table 1). We provide high-level descriptions below and include additional details in Supplementary Section C.

**Table 1:** Statistics of the simulated and biological datasets used in this study. *n* denotes the number of species and *k* denotes the number of single-copy or multi-copy genes.

| Simulated | $n$ | $k$ | $\ S - G\ ^\S$ | $\ G - \hat{G}\ $ | Gene len. (bp) | See |
| --- | --- | --- | --- | --- | --- | --- |
| ILS-only | 101 | 1000 | 30%–58% | 23, 31, 42, 55% | 1600, 800, 400, 200 |  |
| ILS+GDL | 21–1001 | 50–10000 | 15%–78%* | 18% – 56% | 50, 100, 500 | Table S7 |
| ILS+HGT | 51 | 1000 | 30%–68% | 28% | 1000 | Table S8 |
| Biological | $n$ | $k$ | Type | $\ \hat{S} - \hat{G}\ ^\P$ | Total len. (bp)<sup>#</sup> | See |
| Birds | 363 | 63,430 | ILS | 31.1% | 63,430,000 | Table S9 |
| Bees | 32 | 853 | ILS | 39.7% | 576,041 | Table S9 |
| Mammals | 37 | 424 | ILS | 26.6% | 1,385,220 | Table S9 |
| Fungi | 16 | 706<sup>†</sup>, 7180<sup>‡</sup> | ILS+GDL | NA | 286,114 | Table S9 |
| Plants(1kp) | 80 | 424<sup>†</sup>, 9610<sup>‡</sup> | ILS+GDL | 48.0% | 290,718 | Table S9 |
| Eudicots | 40 | 345<sup>†</sup>, 2573<sup>‡</sup> | ILS+GDL | 50.5% | 262,343 | Table S9 |
| AB core | 72 | 49 | HGT | 69.3% | 10,522 | Table S9 |
| AB non-rib. | 108 | 38 | HGT | 61.8% | 6,534 | Table S9 |
| AB WoL | 10,575 | 381 | HGT | 53.9% | 1,162,421,084 | Table S9 |

<sup>§</sup> Average discordance between two types of trees is denoted by $\|a - b\|$ and is measured using the average RF distance. $S$ and $\hat{S}$ denote model and estimated species trees; $G$ and $\hat{G}$ denote true and estimated gene trees.
\* RF between true gene trees and the true locus tree, which is only due to ILS.
<sup>†</sup> single-copy genes, used in concatenation.
<sup>‡</sup> multi-copy genes used for ASTRAL-Pro and CASTLES-Pro.
<sup>¶</sup> RF distance between single-copy gene trees and species trees estimated by ASTRAL or ASTRAL-Pro (NA when not available).
<sup>#</sup> The total length of the sequence alignments for the single-copy loci.

### Simulations

We studied three sets of simulated datasets with gene tree discordance due to ILS, ILS+GDL, and ILS+HGT (Table 1). All simulated datasets are generated using SimPhy (Mallo et al., 2016); however, we modified SimPhy to output model species trees with SU branch lengths using mutation rates already present in SimPhy simulations. Gene sequences were simulated under GTR+Γ model, and gene trees were estimated from these alignments using FastTree-2 (Price et al., 2010). The ILS-only dataset was reused from Tabatabaee et al. (2023) and has gene alignments of length 200bp – 1600bp to control gene tree estimation error (GTEE). The level of ILS is heterogeneous across replicates, with mean equal to 46% according to average Robinson and Foulds (1981) (RF) distance between model species trees and true gene trees (AD for short). The GDL+ILS dataset was reused from Willson et al. (2022, 2023) and has two levels of ILS (low and high), six duplication rates (10^−13^ – 10^−9^), three sequence lengths, various numbers of species, and genes (Table 1). The loss rate relative to the duplication rate is set to 1, 0.5, or 0. For ILS+HGT, we recreated a dataset by Davidson et al. (2015) with 30% AD due to ILS and six levels of HGT rates, leading to up to 68% AD (Table S8). The average number of HGT events per gene for the six model conditions starts from 0 to 0.08, 0.2, 0.8, 8 and 20, corresponding to HGT rates 10^−9^× (0, 2, 5, 20, 200, and 500).

In all simulations, we estimate branch lengths on the fixed true species tree topology. We measure branch length estimation error using three metrics: mean absolute error 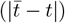, mean logarithmic error 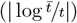, and the bias of the estimated length 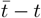 (*t* and 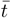 are true and estimated branch lengths, resp.) averaged across all species tree branches. The log error emphasizes short branches, while the absolute error and bias emphasize long branches. We remove the outgroup before measuring the branch length estimation error. For all methods, we replace negative and zero branch lengths with a small pseudo-count (10^−6^) before calculating error metrics.

We compare CASTLES-Pro to CASTLES, ERaBLE, FastME (Lefort et al., 2015) used on matrices of average patristic distances (referred to as FASTME(AVG)), and concatenation using maximum likelihood with RAxML (Stamatakis, 2014). All these methods are only designed to work with single-copy genes; hence, for datasets with GDL, we create a two-step pipeline where we first use the method DISCO (Willson et al., 2022) to decompose gene family trees into single-copy gene trees, which we then pass to branch length estimation methods. These two-step methods are referred to as CASTLES-DISCO, ERaBLE-DISCO, and FastME(AVG)-DISCO. To perform concatenation with multi-copy input, we use the CA-DISCO technique of Willson et al. (2022). Sequences for each gene family are broken up into single-copy loci, and these loci are concatenated into a super-alignment. We use RAxML on this alignment to optimize branch lengths on the fixed true species tree topology. Note that DISCO can produce trees with high levels of missing data, and ERaBLE and FastME(AVG) can fail on inputs with missing data. To enable these methods to run on DISCO output, we imputed the missing values in the distance matrix of each gene tree by the average patristic distances among gene trees that do include the pair of taxa associated with the missing value. Finally, while the species tree estimation method SpeciesRax (Morel et al., 2022) can produce branch lengths in substitution units, we did not include it in this study as it cannot estimate branch lengths on a fixed input topology.

### Biological datasets

We reexamined nine biological datasets with different sources of gene tree discordance (Table 1, Table S9). As examples of datasets with ILS, we analyzed the birds dataset by Stiller et al. (2024), bees by Bossert et al. (2021), and mammals by Song et al. (2012). For GDL, we analyzed two plant datasets (Chanderbali et al., 2022; Wickett et al., 2014) and a fungal dataset (Butler et al., 2009) that included multi-copy gene family trees. Finally, following Moody et al. (2022a), we analyzed three bacterial datasets (Petitjean et al., 2015; Williams et al., 2020; Zhu et al., 2019) as examples of cases with high rates of HGT, with a focus on the length of the branch that separates archaea and bacteria at the root of the tree of life (AB branch).

We compare the branch lengths produced by CASTLES-Pro on ASTRAL or ASTRAL-Pro topologies to concatenation branch lengths drawn on either ASTRAL or concatenation topologies. For single-copy datasets, we used the ASTRAL topology for both concatenation and CASTLES-Pro branch lengths. For the three datasets with GDL, since directly using concatenation on multi-copy gene sequences was not possible, we compared the concatenation topology on single-copy genes from original studies to ASTRAL-Pro topology furnished with CASTLES-Pro branch lengths run on multi-copy genes (the two trees differed by 8–9% RF). In these cases, we focus on branches that are shared between the two trees. For birds, bees, and mammals, ASTRAL trees were already available, while for fungi, we inferred a tree using ASTRAL-Pro2 (Zhang and Mirarab, 2022a) using all multi-copy gene trees, and for small bacterial datasets, we inferred a tree using ASTRAL-III (Zhang et al., 2018). For 1KP, single-copy concatenation and multi-copy ASTRAL-Pro trees have 80 taxa in common, which we use. For some datasets, original studies ran concatenation on a subset of sites from the sequence alignments: For the fungi dataset, 30,000 sites were sampled from 706 orthologs, and for the WoL bacterial dataset, 100 sites were randomly selected from sites with less than 50% gaps for each of the 381 marker genes.

## Results

### ILS-only simulations

CASTLES-Pro has the best accuracy across all GTEE levels of this dataset, followed by CASTLES (Figs. 2 and S6). Distance-based methods come next, with TCMM outperforming ERaBLE and FastME(AVG), and concatenation is the least accurate overall. Relative accuracy of methods is mostly consistent across different alignment lengths (Fig. 2A) and levels of ILS (Fig. 2B). As alignment length increases (and GTEE decreases), concatenation remains stable while CASTLES-Pro and distance-based methods become successively better. Improvements of CASTLES-Pro are mostly due to better terminal branches, which are substantially more accurate than the other methods in all conditions (Fig. 2). On internal branches, the relative accuracy depends on the condition; CASTLES-Pro is better for true gene trees and slightly worse than the distance-based methods and CASTLES in the highest GTEE level. Finally, note that pairing TCMM with CASTLES-Pro, as described by Arasti et al. (2024), substantially reduces the error of TCMM, but in these ILS-only conditions, the combination is not as good as CASTLES-Pro alone (Fig. S6).

**Figure 2:**
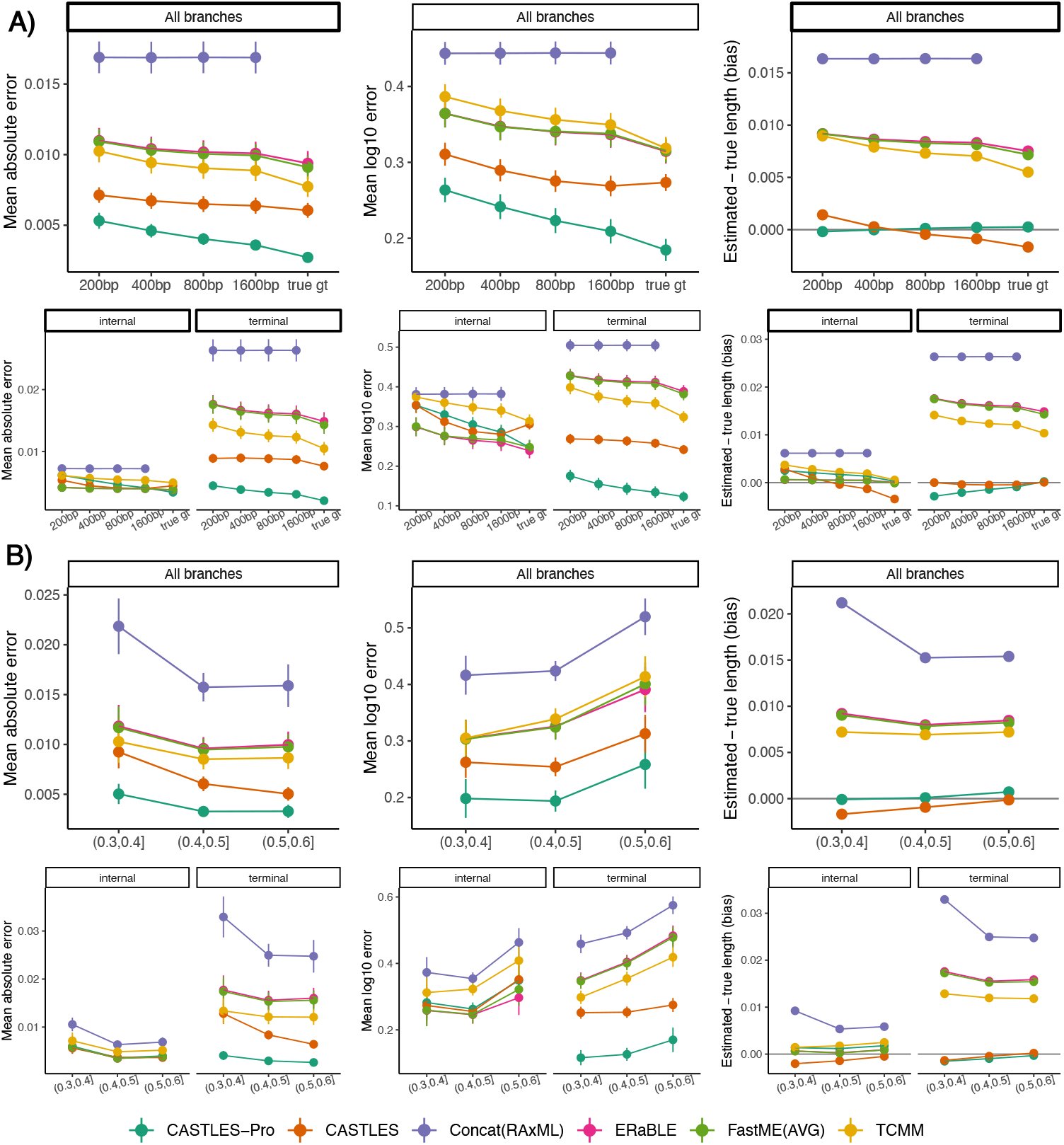
Mean absolute error, mean log error and bias for different branch length estimation methods on 100-taxon simulated ILS datasets. The average ILS level on this dataset is 47% AD. The number of genes is 1000, and the number of replicates is 50. A) Varying gene tree error (GTEE) (*x*-axis) where the GTEE level changes between 0% for true gene trees to 55% for gene trees estimated from 200bp alignments. B) Varying the level of ILS (x-axis) for conditions with 1600bp sequence length. The number of replicates in the three ILS bins are 9, 29, and 12, respectively. See also Fig. S6 for comparison between CASTLES-Pro and CASTLES-Pro+TCMM.

In addition to better accuracy, CASTLES-Pro has the lowest overall bias (Figs. 2 and S6). In particular, concatenation has a substantial overestimation bias for terminal branches, but it also overestimates internal branches to a lesser degree. CASTLES shifts from a small underestimation bias for true gene trees to a minor overestimation bias for gene trees with high GTEE; this is due to the effects of the imprecise Lambert approximation used in CASTLES, which is fixed in CASTLES-Pro. Distance-based methods also exhibit an overestimation bias, though it is smaller than that of concatenation. Evaluating terminal and internal branches separately (Fig. 2) shows that CASTLES-Pro is unbiased for both terminal and internal branches when given true gene trees; as the GTEE increases, it suffers a small overestimation bias for internal branches and a similar underestimation bias for terminal ones; thus, effects of gene tree error are reduced but not fully eliminated in CASTLES-Pro. Finally, distance-based methods have a small overestimation bias for internal branches that increases as GTEE increases, and a much larger bias for terminal branches.

The trends for log error, which emphasizes short branches more than absolute error, are similar, and CASTLES-Pro is the most accurate method overall, followed by CASTLES, distance-based methods, and finally concatenation. The only difference, according to log error, is that TCMM is overall slightly less accurate than the other two distance-based methods (ERaBLE and FastME(AVG)) but still more accurate than them for terminal branches.

### GDL+ILS simulations

In the presence of GDL and ILS, CASTLES-Pro has the lowest error and bias in many but not all conditions, with concatenation used with DISCO (CA-DISCO) performing better with low ILS according to some metrics (Fig. 3). CASTLES-Pro is always the most accurate method in the high ILS conditions, followed by CADISCO, but the gap is particularly large for lower duplication rates (Fig. S7) or large numbers of gene trees (Figs. S8 and S9). For the low ILS condition, CASTLES-Pro is still better than CA-DISCO according to log error in most conditions but is outperformed in some conditions according to the mean absolute error metric; in particular, CA-DISCO clearly outperforms CASTLES-Pro with 20 species according to the mean absolute error metric regardless of the duplication rates (Fig. S7), number of genes (Fig. S8), or sequence length (Fig. S10). However, with more species, CASTLES-Pro either outperforms or matches CA-DISCO even with low ILS (Figs. 3 and S11). Since log error emphasizes short branches more than mean absolute error, these trends suggest that CASTLES-Pro is doing a consistently better job at estimating short branches, whereas concatenation is sometimes better at estimating long branches. Other methods are less competitive. CASTLES run on DISCO decomposed gene trees is less accurate than CASTLES-Pro in most conditions across both 20-taxon and 100-taxon datasets (Figs. S7 to S12). Distance-based methods are the least accurate in almost all conditions.

**Figure 3:**
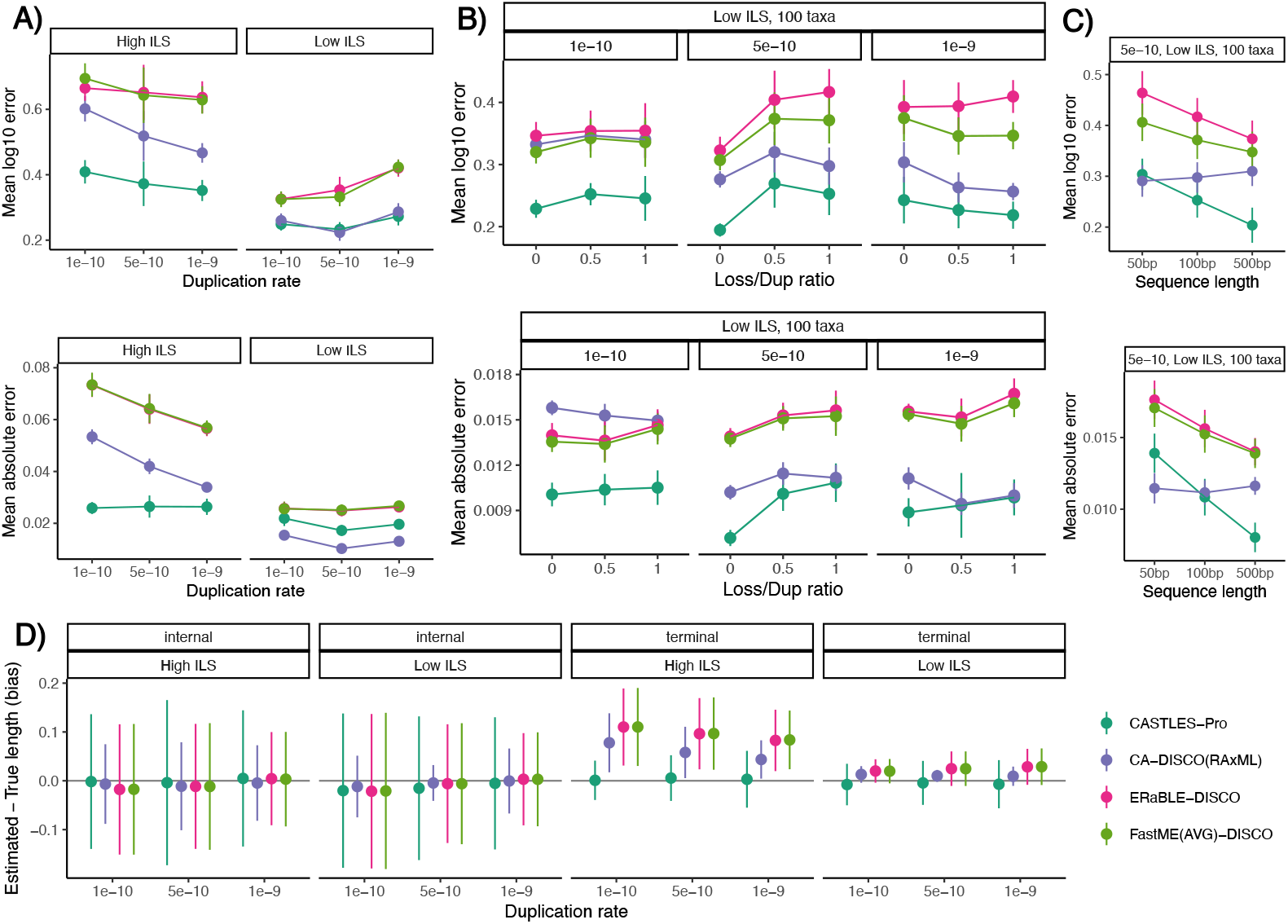
Mean log error, mean absolute error, and bias of branch lengths for simulated GDL+ILS datasets, varying number of species, duplication rate, ILS level, and sequence length. When not specified, each parameter is set to default: 1000 genes, 100bp sequence length, equal loss and duplication rates, 20 species. A) 20-taxon datasets varying duplication rate and ILS rate. B) 100-taxon datasets, low ILS condition, varying duplication rate and loss/dup ratio. C) 100-taxon datasets, 5e-10 duplication rate, low ILS condition, varying sequence length. D) Bias divided by terminal and internal length for the conditions in panel A. The number of replicates is 10. See Fig. S7-S12 for full results.

The number of genes, sites per gene, and species all impact accuracy in various ways. As the number of genes increases, the error of CASTLES-Pro drops faster than CA-DISCO for both 20 (Fig. S8) and 100 species (Fig. S9). Similarly, increasing the sequence length and thus decreasing GTEE does not impact CA-DISCO, but makes CASTLES-Pro and other methods more accurate (Fig. S10), with CASTLES-Pro improving the fastest, especially with 100 species (Fig. 3C). The number of species has a mixed impact, which depends on the rate of duplication, the measure of error, and the choice of method (note that tree heights are fixed when the number of species changes, creating shorter branches with more species). The impact of the duplication rate also depends on the level of ILS and method. Overall, CASTLES-Pro is relatively robust, retaining similar error and bias levels across different duplication rates (Figs. 3 and S7). Methods that rely on DISCO to decompose the trees tend to become better with higher duplication rates, especially with high ILS. Increasing loss rates, however, can increase the error of CASTLES-Pro in some cases but does not introduce any discernable bias (Figs. 3 and S12).

Overall, CASTLES-Pro has a lower bias than other methods, especially for terminal branches (Fig. 3D, Fig. S7 to Fig. S12). CA-DISCO clearly overestimates terminal branches, especially for higher ILS levels; in contrast, CASTLES-Pro does not have a clear bias for terminal branches. For internal branches, all methods are less biased, with CA-DISCO and CASTLES-Pro performing slightly better for low and high ILS conditions, respectively. The distance-based methods also have a clear overestimation bias for terminal branches. Overall, the most glaring form of bias is for terminal branches for high ILS conditions in all experiments, a problem that CASTLES-Pro eliminates.

### Scalability

CASTLES-Pro also has a runtime advantage on the 100-taxon GDL datasets, and its advantage becomes more clear as the number of genes increases (Fig. S13). In particular, for 10,000 genes, CASTLES-Pro takes on average less than 1 minute to estimate branch lengths on a fixed tree topology, while CA-DISCO takes about 124 minutes (Table 2). The gap between CASTLES-Pro and CA-DISCO widens as gene family trees become larger (Table S7); with the highest duplication rate (10^−9^) and no loss, CASTLES-Pro finishes in 4 minutes on average, while CA-DISCO takes more than 25 hours on average (Fig. S13). In terms of memory usage, CASTLES-DISCO and CASTLES-Pro are almost identical and use much less memory than other methods (Table 2 and Fig. S13). Finally, in the model condition with 1000-taxon trees and 1000 genes, CADISCO and distance-based methods fail due to memory limit given 128GB of RAM, while CASTLES-Pro and CASTLES-DISCO finish in 2 and 11 minutes on average resp. and use less than 4 GB of memory.

**Table 2:** Runtime and peak memory usage of different methods for 100-taxon GDL+ILS dataset for 10,000 genes with sequence length of 100bp. The duplication rate is 5 *×* 10^−10^ with equal loss rate. The results are averaged across 10 replicates. The runtime does not include gene tree estimation or species tree topology estimation time, as all methods draw branch lengths on a fixed tree topology. See also Fig. S13.

|  | time (minutes) | peak memory (GB) |
| --- | --- | --- |
| CASTLES-Pro | 0.96 | 4.09 |
| CASTLES-DISCO | 6.45 | 3.64 |
| CA-DISCO(RAxML) | 123.72 | 29.26 |
| ERaBLE-DISCO | 43.35 | 12.78 |
| FastME(AVG)-DISCO | 19.61 | 8.81 |

### HGT+ILS simulations

The first four model conditions of this dataset have low HGT rates and little discordance beyond ILS (increasing from 30% with ILS-only to 34%). Therefore, the log error for all methods has almost no change across these four conditions, and the mean absolute error fluctuates within the bounds of standard error (Fig. S14A). However, the last two conditions have substantially higher HGT rates, with 53.4% and 68.4% total discordance (Table S8). Comparing the last three conditions, both log and mean absolute error for all methods generally increase for higher HGT rates. Distance-based methods are generally the least accurate across different conditions, except for the highest HGT rate, where concatenation has a higher mean absolute error. CASTLES-Pro is the most accurate for both metrics across all conditions, except for the highest HGT rate, where it has a tie with CASTLES in terms of log error. While TCMM is inaccurate, following CASTLES-Pro by regularized TCMM, as detailed by Arasti et al. (2024) further improves its accuracy and obtains the best results overall. The gap between CASTLES-Pro (with or without TCMM) and concatenation widens as HGT increases in terms of absolute error (i.e., focusing on long branches) but closes for mean log error (focusing on short branches).

In terms of bias, terminal and internal branches again show different patterns (Fig. S14B). CASTLES-Pro and CASTLES have an underestimation bias in all model conditions, especially for internal branches. This underestimation mostly disappears if CASTLES-Pro is followed by TCMM. Concatenation and, to a smaller degree, distance-based methods have a large overestimation bias for terminal branches that increases with HGT rates. The over-estimation of terminal branches by concatenation is, on average, 2.77 times larger than the underestimation for terminal or internal branches by CASTLES-Pro and 5.45 times larger than the underestimation bias of CASTLES-Pro+TCMM over all branches.

### Biological datasets

#### ILS

On the three datasets where biological discordance is likely dominated by ILS (birds, bees, and mammals), we observe that CASTLES-Pro produces shorter lengths than concatenation, especially for terminal branches (Fig. 4). This pattern is more extreme for the birds and bees datasets (13.8% and 10.6% increase in average root-to-tip distance respectively), which have a particularly high level of observed gene tree discordance. In addition, changes are more pronounced for terminal branches (Fig. 4) and especially short terminal branches (Figs. S15 and S16), as expected by theory and results of the simulations.

**Figure 4:**
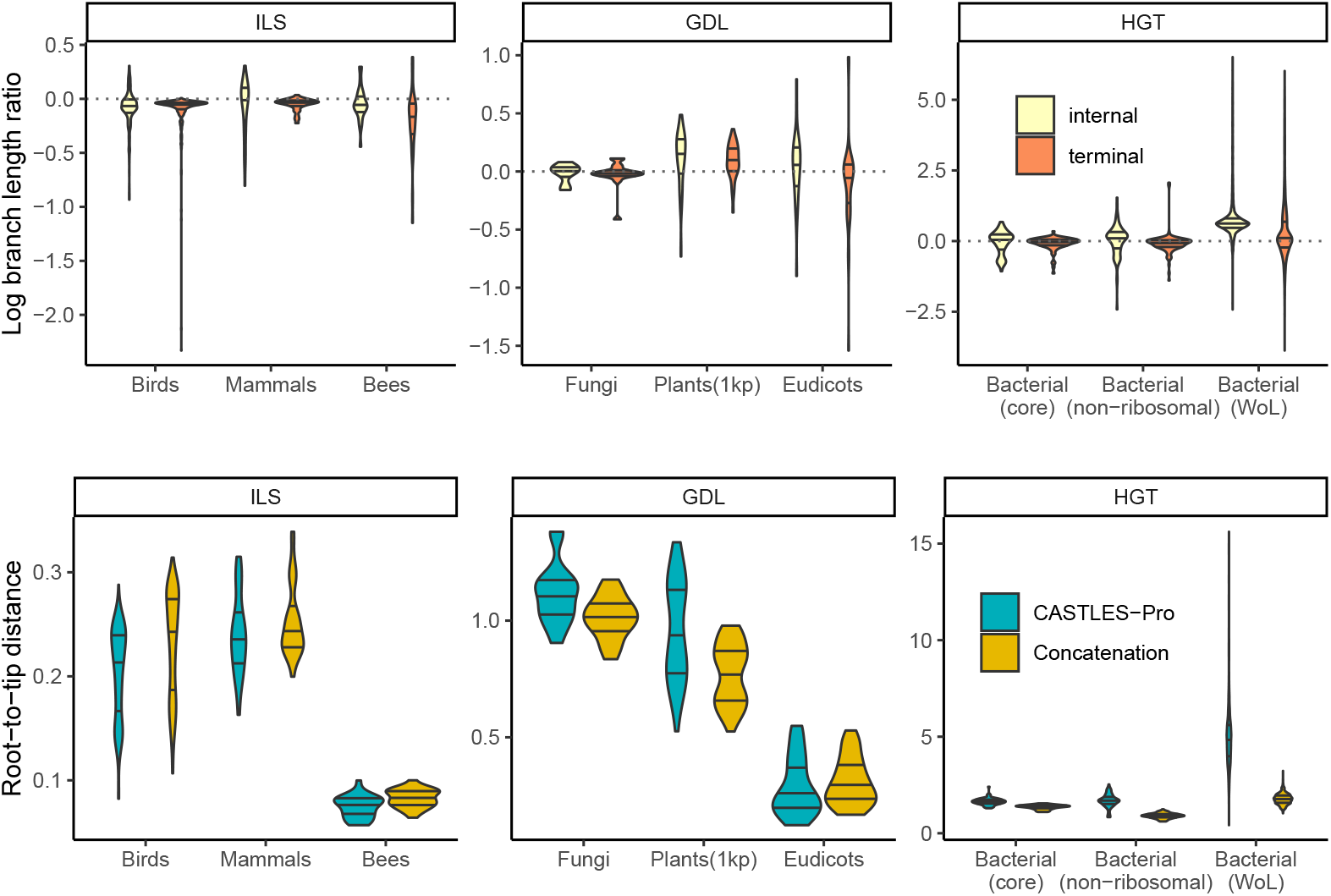
(top) The branch lengths produced by CASTLES-Pro divided by branch lengths of concatenation on nine biological datasets with different sources of gene tree heterogeneity in log scale. (bottom) Distribution of the root-to-tip distance for the CASTLES-Pro and concatenation trees on the nine biological datasets. See also Fig. S24.

For terminal branches of the bird dataset, concatenation has a slightly shorter length for only one species and substantially longer for others. The median (1st and 3rd quantiles) reduction from concatenation to CASTLES-Pro lengths is 0.00333 SU (0.00247, 0.00454); if we assume a mean substitution rate of 0.0026 per million years as estimated by Stiller et al. (2024) and 6.6 years per generation (median across species reported by Houde et al. (2020)), the gap between concatenation and CASTLES-Pro corresponds to the expected 1 CU if *N*_*e*_ = 0.00333*/*0.0026 × 10^6^*/*6.6 = 194*k* (144*k*, 265*k*), which matches previous estimates using PSMC (Nadachowska-Brzyska et al., 2015).

Similarly, for bees, there are only three shorter terminal branches in concatenation compared to CASTLES-Pro, and one of them (*S. schubotzi*) becomes substantially longer. This increase is due to one gene tree with the clearly incorrect terminal length of 2.63 for *S. schubotzi*. Removing this single gene tree or using TreeShrink (Mai and Mirarab, 2018) to filter abnormally long branches both reduce the length of this branch in the CASTLES-Pro output from 0.0454 to 0.0145 or 0.0157, which are below concatenation (Fig. S15). Using original gene trees (including outliers), we see the expected decrease in terminal lengths, with a median of 0.00255 (0.00159, 0.0047), which, divided by the spontaneous mutation rate of 3.5 × 10^−9^ per generation given by Wallberg et al. (2015), corresponds to *N*_*e*_ = 729*k* (452*k*, 1, 350*k*) for 1 CU, which is in line with estimates based on PSMC (Lozier et al., 2023).

Finally, for mammals, four terminal branches become longer in CASTLES-Pro (including one with mislabeled taxa) while others become shorter. The elongated four are due to outliers as using TreeShrink shortens all these four branches, with two becoming shorter than concatenation and the other two remaining only 0.5% and 1.4% longer than concatenation. Even without TreeShrink, terminal branches shrink in CASTLES-Pro by a median of 0.0036 (0.0014, 0.0052), which assuming a per generation mutation rate of 2.5 ×10^−8^ (Pfeifer, 2020) would correspond to 1 CU for *N*_*e*_ = 145*k* (57*k*, 206*k*).

#### GDL

Patterns on GDL datasets differ from ILS-dominated datasets (Fig. 4) perhaps because here concatenation is run on single-copy genes while CASTLES-Pro is run on the full set of multi-copy gene trees. On the 1KP plant dataset, CASTLES-Pro run on 9610 multi-copy gene trees has longer terminal and internal branch lengths than concatenation run on 424 single-copy genes (Figs. 4 and S19), leading to 24.2% higher mean root-to-tip distance. Similarly, on the fungal dataset, CASTLES-Pro based on 7,180 multi-copy gene family trees produces longer branches and 10.1% higher average root-to-tip distance compared to concatenation on 706 single-copy genes (Figs. 4 and S20). On the small and less diverse 40-taxon eudicots dataset, CASTLES-Pro based on 2,573 multi-copy gene trees results in shorter terminal branches but longer internal branch lengths than concatenation based on 345 single-copy genes (Figs. 4 and S18), with 7.7% decrease in average root-to-tip distance. Thus, patterns were different between these two plant datasets and patterns were unlike ILS datasets; we return to this point in the discussions.

#### Micorbial data and AB branch

A long-standing hypothesis has been that bacteria and archaea domains are separated by a long branch (Cox et al., 2008; Gogarten et al., 1989; Iwabe et al., 1989). In contrast, Zhu et al. (2019) (which included some of us) estimated a far shorter length for the AB branch than what was previously reported using a concatenation of 381 marker genes for the branch length estimation step. Moody et al. (2022a) further studied this and other microbial datasets and suggested (among other criticisms) that concatenation can severely underestimate branch lengths on datasets with high levels of HGT, resulting in underestimation of the AB branch length. They report estimates of AB branch as long as 3.3 based on the core gene set of Williams et al. (2020) and 2.52 using 27 most vertically evolving genes selected from a set of manually curated marker genes including both ribosomal and non-ribosomal proteins. We reexamine some of these datasets using CASTLES-Pro.

On all three microbial datasets, we observe a general increase in the length of the internal branches, particularly longer branches, and a small decrease in the length of terminal branches (particularly short ones) for CASTLES-Pro compared to concatenation (Figs. 4 and S23). Since internal branches increase more than terminal branches decrease, we observe a substantial increase in the average root-to-tip distance on all three datasets (Figs. 4, 5 and S21 to S23). For the WoL dataset, branches change dramatically between the methods; however, note that Zhu et al. (2019) limited itself to 100 sites per gene due to scalability limitations of concatenation, while CASTLES-Pro uses gene trees estimated from full-length sequence alignments. The concatenation tree has a long tail of branches with 1e-6 or 2e-6 length, corresponding to no-event branches among 100 × 381 sites chosen, but no such tail exists for CASTLES-Pro (Fig. S23) since it uses all sites.

**Figure 5:**
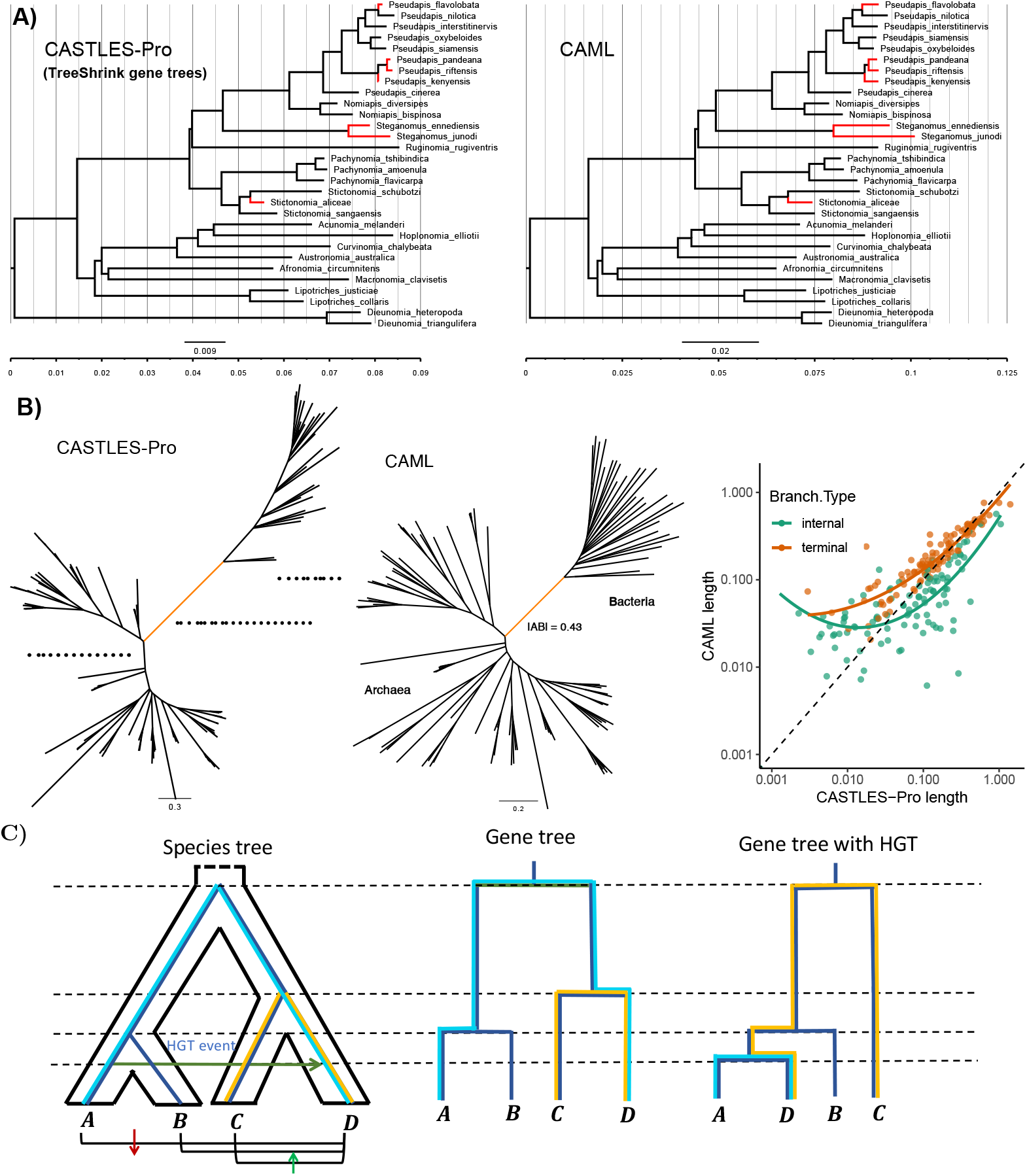
A) Comparison between the branch lengths of CASTLES-Pro (with TreeShrink gene trees) and CAML on the 32-taxon 853-gene bees dataset of Bossert et al. (2021) after removing the outgroup taxa *Lasioglossum albipes* and *Dufourea novaeanglia*. We used the ASTRAL topology from the original study, and used concatenation and CASTLES-Pro to draw branch lengths on this topology. The branch lengths that are at least 2x shorter in CASTLES-Pro tree compared to the concatenation tree are highlighted in red. B) Comparison between the branch lengths produced by CASTLES-Pro and concatenation on the 108-taxon bacterial dataset with 38 non-ribosomal genes. The branch highlighted in orange separates domain Archaea from Bacteria (AB branch). C) HGT, if ignored (as in concatenation) can make branches longer or shorter; the HGT event shown by the dark green arrow creates the gene tree shown on the right, which has a shorter distance from A and B to D, but higher distances from D to C. This reduces branch length for some branches in the species tree (blue) and increases it for others (yellow). See also Figs. S15-S23.

On all three datasets, CASTLES-Pro substantially increased the AB length compared to concatenation (Table 3) and made them closer to those estimated by Moody et al. (2022a). The increases are dramatic (17×) on the WoL dataset which contains highly discordant genes, substantial (2.5×) on the less discordant non-ribosomal genes, and relatively small (1.3×) on the presumably HGT-free core genes. On the two small less discordant datasets, there are orders of magnitude more matching quartets than non-matching quartets, signifying lack of discordance, and in both cases, length of matching quartets is much longer than non-matching ones (Table 3). In contrast, on WoL, the number of matching and non-matching quartets are both very large, and matching quartets are only 1.7x more than non-matching ones. Nevertheless, the length of the AB branch in all three cases remains close to the average AB length in the matching quartets. Overall, CASTLES-Pro produces longer internal branches and longer AB lengths for all three bacterial datasets compared to concatenation, a trend that agrees with the observations of Moody et al. (2022a), who suggest that concatenation can underestimate branch length in the face of high HGT.

**Table 3:** AB branch length on the three bacterial datasets estimated by CASTLES-Pro, TCMM and concatenation. 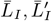 refer to the average AB branch length in matching and non-matching gene trees, respectively, and *C*_*M*_, *C*_*NM*_ refer to the number of matching and non-matching quartets around the AB branch. *d* refers to the length of the AB branch in coalescent units.

| | $\bar{L}_I$ | $\bar{L}'_I$ | $C_M$ | $C_{NM}$ | $d$ | CASTLES-Pro | TCMM | CAML |
| --- | --- | --- | --- | --- | --- | --- | --- | --- |
| Core | 1.92 | 0.01 | 922,226 | 32 | 9.83 | 1.79 | 1.94 | 1.43 |
| Non-ribosomal | 1.06 | 0.08 | 1,270,289 | 11,624 | 4.30 | 1.05 | 0.87 | 0.43 |
| WoL | 2.74 | 0.74 | 750,795,570,342 | 444,568,398,602 | 0.58 | 2.68 | 1.89 | 0.16 |

## Discussion and future work

We introduced CASTLES-Pro, a summary method that can furnish a given species tree with substitution-unit branch lengths accounting for GDL and ILS based on a given set of potentially multi-copy gene trees. Our simulations with ILS alone or ILS+GDL showed that CASTLES-Pro is more accurate than other methods in most conditions and is also more scalable, easily running on datasets with tens of thousands of species or genes (Table S10) in less than an hour. In the face of HGT, which CASTLES-Pro does not directly model, it was still more accurate than concatenation but had room for improvement. In particular, using CASTLES-Pro with TCMM of Arasti et al. (2024) reduced its bias for high HGT cases. Paired with summary methods that infer the topology, this advance makes the two-step approach to species tree inference far more useful than before. The ASTER package outputs trees with SU branch lengths when used with ASTRAL-IV (for single-copy genes) or ASTRAL-Pro-2 (for multi-copy genes) using the CASTLES-Pro algorithm. These trees can readily be used as input to downstream analyses such as dating.

The negative impact of concatenation on branch lengths depended on the cause of discordance, as biological analyses clearly show (Fig. 4). For ILS, as expected, terminal branch lengths, especially shorter ones, are over-estimated using concatenation, but internal branches have far less bias overall. This imbalance between the error for terminal and internal branches can lead to unexplainable patterns in downstream analyses, such as diversification rates. Note that for internal branches, depending on the coalescent length of surrounding branches, we may still have overestimation or underestimation using concatenation, but looking across all internal branches, those effects diminish.

In contrast to ILS, on GDL datasets where CASTLES-Pro was given all loci and sites per locus and concatenation was based on fewer genes and sites, CASTLES-Pro generally produced longer branches. One explanation for the increase instead of decrease in branch lengths in GDL datasets is ascertainment bias. The small portion of loci that happened to be single-copy tend to be the most conserved ones, giving a biased picture of the substitution rates. By allowing the use of all multi-copy gene trees, CASTLES-Pro reveals higher genome-wide substitution rates compared to single-copy genes. An alternative explanation is that single-copy gene trees are easier to align due to being smaller (and more conserved), and the higher branch lengths from CASTLES-Pro could be a result of over-alignment in multi-copy genes or perhaps saturation. The real GDL data likely suffer from a mixture of both, especially given that angiosperm data (which span a far shorter evolutionary time) did not experience branch length increase.

For HGT, our simulations showed an over-estimation bias for concatenation, whereas, on real microbial data, we seemed to observe the opposite. For the AB branch (but also others), concatenation under-estimates lengths compared to CASTLES-Pro and compared to using HGT-free genes. The difference is likely due to the type of HGT events. Any single HGT event both increases and decreases divergence for some pairs of taxa compared to the species divergence, leading to both under-estimation and over-estimation bias for different branches (see Fig. 5C). Our simulations using Simphy augment ILS with *random* HGT events, each of which can create bias in either direction for individual branches. When many such events accumulate, they can cancel each other out, leading to a less clear HGT signature and leaving us with impacts of ILS. Real biological data often experience highways of HGT when large numbers of genes are transferred between two points of the tree. Such highways are expected to have happened around the AB branch (Beiko et al., 2005; Puigb`o et al., 2009). One expects HGT highways between two branches to create a strong under-estimation bias for branches connecting the donor and recipient; a large portion of the concatenated alignment will have reduced divergence between pairs of species, one from the donor and one from the recipient (see Fig. 5C). Our results are consistent with the claim by Moody et al. (2022a) that the AB length seems to be underestimated by concatenation for this reason.

The underestimation around the AB branch seemed to be fixed in CASTLES-Pro, but the reason needs explanation since this phenomenon is separate from the coalescent dynamics CASTLES-Pro models. When a branch has a long estimated CU length, the relatively few gene tree quartets that disagree with the species tree can have abnormally long lengths, even exceeding the matching ones on average. This is exactly the situation for the AB branch (Table 3). In such cases, CASTLES-Pro resorts to using the mean length of matching gene tree quartets for the species tree and thus effectively ignores coalescent effects, which is defensible for a branch with a large CU length. In doing so, it eliminates gene tree quartets that disagree with the species tree, which are presumably due to HGT for the AB branch. This feature (and not expectations under coalescence) leads CASTLES-Pro to output a large distance for the AB branch in contrast to concatenation.

Our method paves the way for a new four-stage phylogenomics pipeline that uses scalable methods in each step: Estimate gene trees independently, estimate the species tree topology by summarizing the gene tree topologies, estimate the branch lengths using CASTLES-Pro (optionally followed by TCMM for high HGT), and date the tree using scalable methods such as TreePL (Smith and O’Meara, 2012) or MD-CAT (Mai et al., 2024). With this pipeline, we can easily handle datasets with thousands of species and thousands of genes.

## Acknowledgment

This work is supported by the National Institute of Health (1R35GM142725). We thank Tandy Warnow for valuable input, including the suggestion to use DISCO with concatenation and other tools, and Edward L. Braun for helpful comments and pointers.

## Supplementary Materials

### A. Details of the Recursive Algorithm

In this section, we will first provide an *O*(*n*^2^) algorithm for computing branch lengths for all internal and terminal branches. Notice, the complexity of this algorithm can be improved to *O*(*nH* log *n*), where *H* denotes the average height of gene family trees, using the dynamic programming algorithm in CASTLES. For conciseness, we use *A, B, C, D* to denote sets of taxa and use *a, b, c, d* to denote individual taxa. Let 𝒢 be the set of gene trees.

To compute all branch lengths, it is sufficient to compute the following counters for a set of ordered leafset quadripartitions, in which each leafset quadripartition (*A, B, C, D*) – up to permutations – corresponds to an internal branch:

- *n*(*A, B*; *C, D*): the number of quartet and gene tree combinations (*a, b, c, d, G*) ∈ *A* × *B* × *C* × *D* × 𝒢 such that *G* ↾ {*a, b, c, d*} has topology *ab*|*cd*.
- *x*(*A, B*; *C, D*): the total internal branch lengths of quartet trees in the form of *G* ↾ {*a, b, c, d*} with topology *ab*|*cd*, where (*a, b, c, d, G*) ∈ *A* × *B* × *C* × *D* × 𝒢.
- *a*(*A*; *B*; *C, D*): the total length of the terminal branches leading to *A* in quartet trees in the form of *G* ↾ {*a, b, c, d*} with topology *ab*|*cd*, where (*a, b, c, d, G*) ∈ *A* × *B* × *C* × *D* × 𝒢.

All three counters for each quadripartition (*A, B, C, D*) can be computed in a single post-order traversal of the gene tree nodes in *O*(*n*) using Algorithm S1. Therefore, computing counters for all *O*(*n*) quadripartitions has time complexity *O*(*n*^2^). Notice that Algorithm S1 assumes that all input gene trees are fully resolved, consistent with the presumption of ASTRAL-Pro.

#### Algorithm S1

Recursive algorithm. The input is a set of gene trees 𝒢 and an ordered quadripartition of the leafset (*A, B, C, D*), and the output are *n*(*A, B*; *C, D*), *x*(*A, B*; *C, D*), and *a*(*A*; *B*; *C, D*). For each node *u* we keep a list of counters 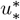 described in Figs. S1 and S2.

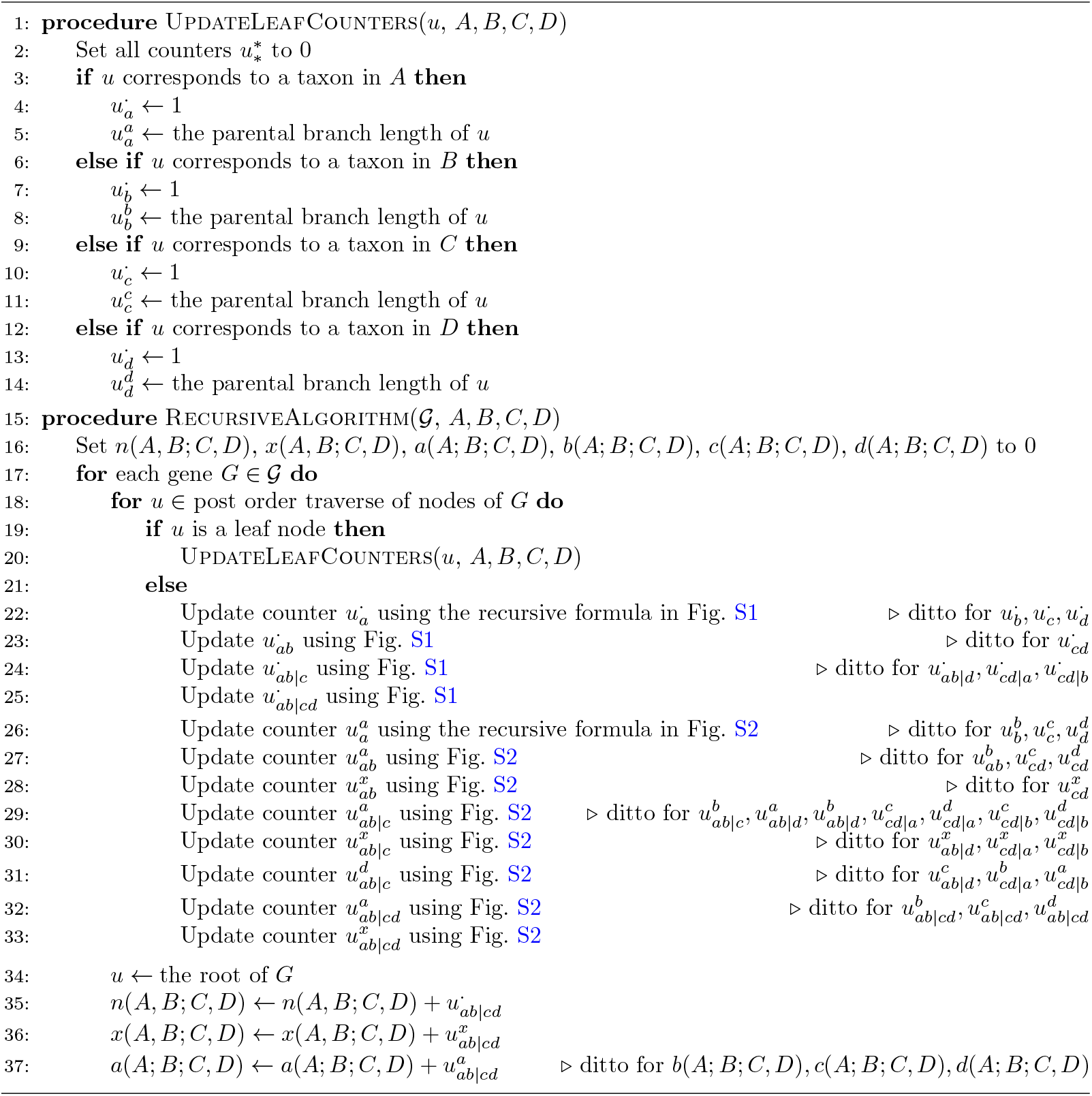

**Figure S1:**
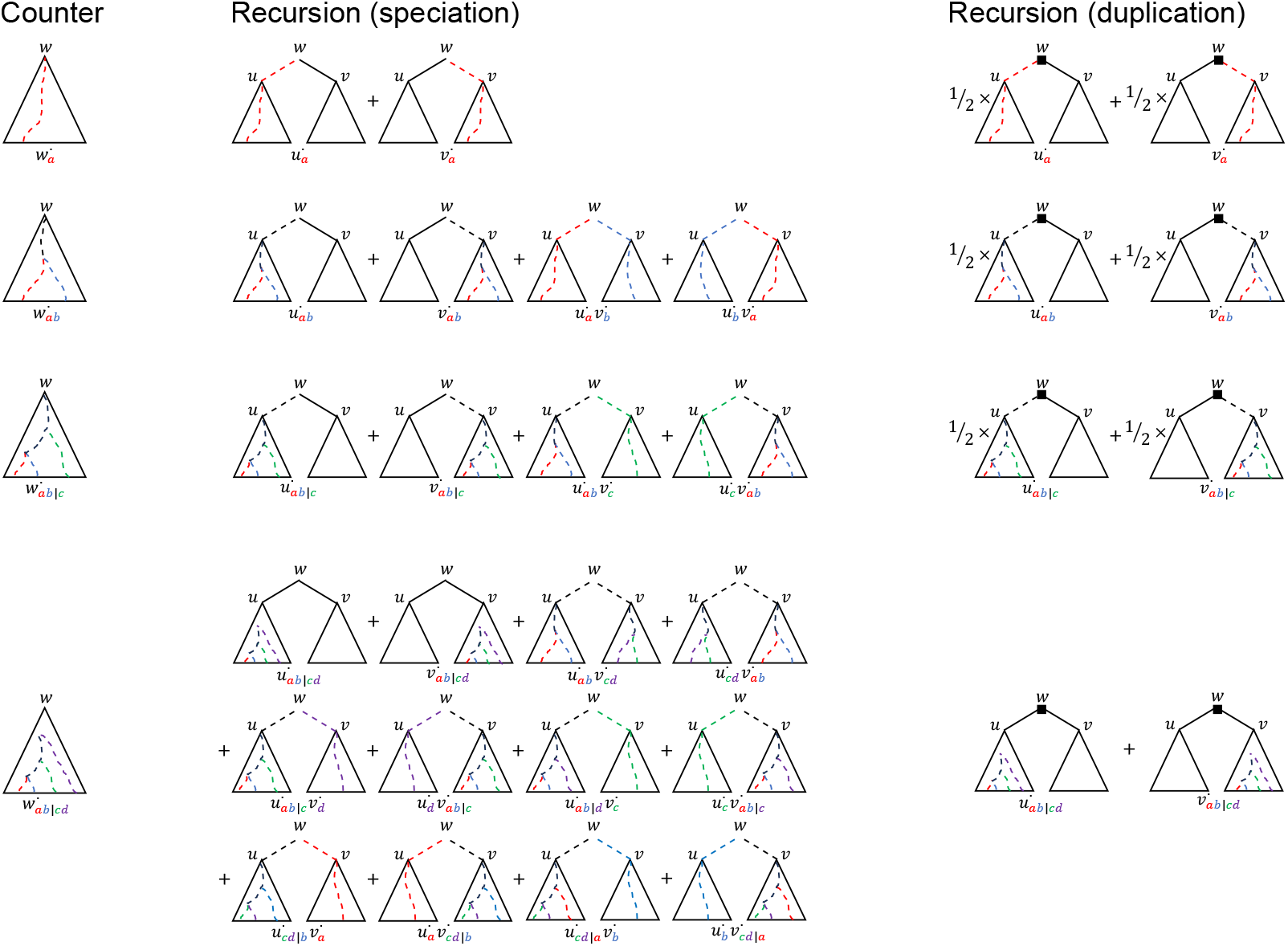
Additional counters for computing the weighted count of gene tree quartets aligning with the species quadripartition *ab*|*cd*. To the left presents a list of counters for each internal node *w*; in the middle illustrates how to recursively compute these counters from counters of the two children of *w* when *w* corresponds to a speciation event; to the right illustrates how to compute these counters when *w* corresponds to a duplication event. For example, 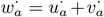 if *w* corresponds to a speciation event, and 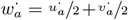 if *w* corresponds to a speciation event (*u* and *v* denote the children of *w*). The total weighted count of gene tree quartets aligning with the species quadripartition *ab*|*cd* is calculated as 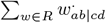, with *R* representing the set of root nodes across all gene trees.

**Figure S2:**
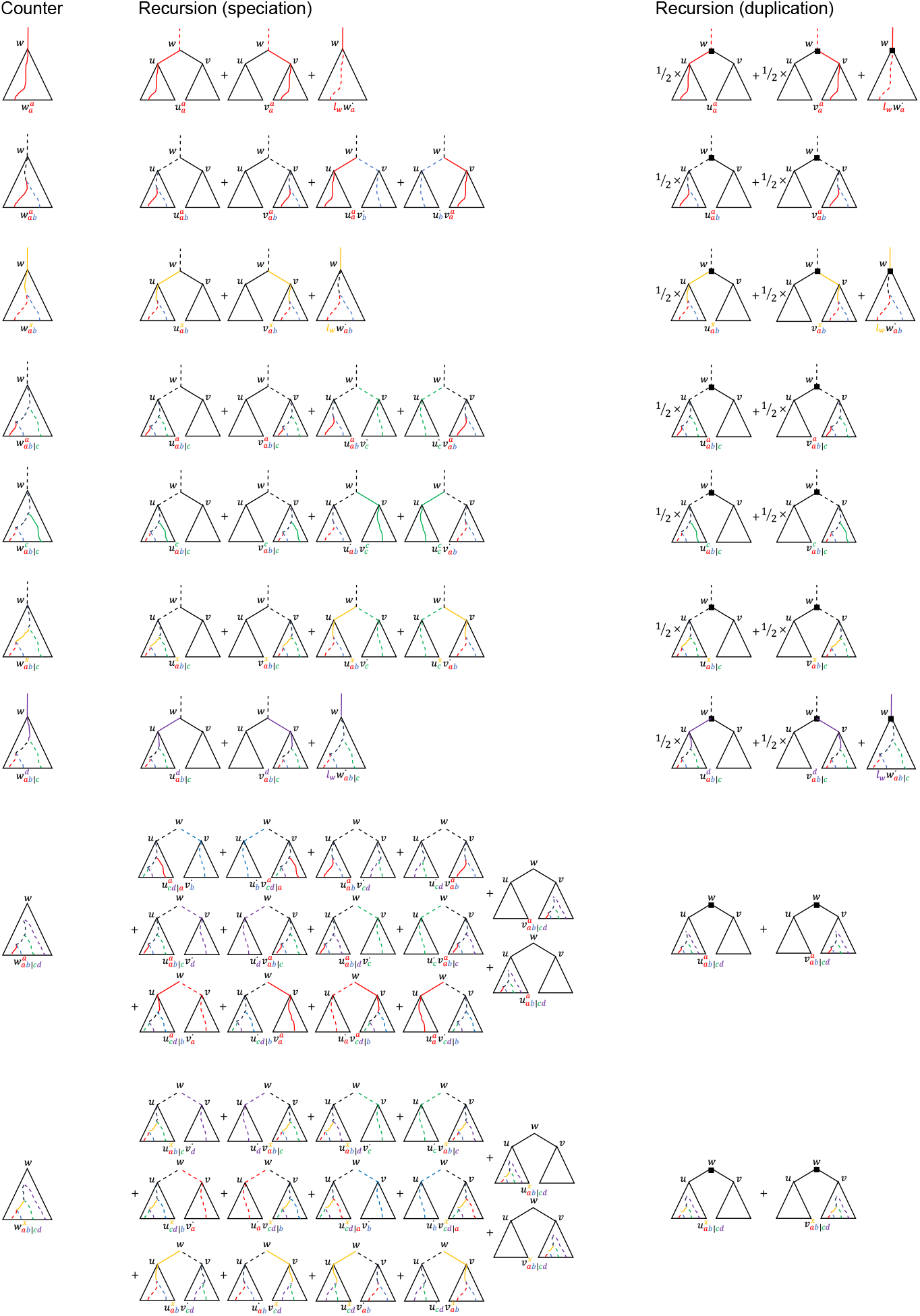
Counters for computing the weighted sums for internal and terminal branch lengths of gene tree quartets aligning with the species quadripartition *ab*|*cd*. The weighted mean internal branch lengths related to *ab*|*cd* are 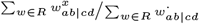 for the matching case and 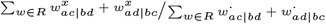 for the non-matching cases (similarly computed by permuting *a, b, c*, and *d*). To compute the weighted mean for terminal branch lengths for the matching case, we use 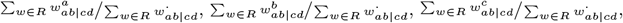, and 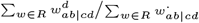, respectively; to compute the weighted mean for terminal branch lengths for non-matching cases, we use 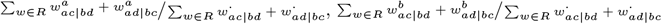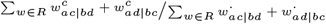, and 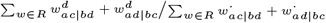, respectively.

### B. CASTLES-Pro’s Equations

Tables S4-S6 summarize the exact and simplified formulas for expected branch lengths in matching and non-matching gene trees under the MSC, and final formulas for estimating branch lengths of an unbalanced or balanced quartet species tree that were used in CASTLES (Tabatabaee et al., 2023); in both figures, parameters are named according to Fig. S3. CASTLES-Pro uses much of the same formulas, but computes them in a different way that leads to improvements in accuracy. The changes for the internal branches (avoiding the Taylor approximation and ILS-aware weighting) are described in the main text, and here we describe the changes in calculating the terminal branches.

**Figure S3:**
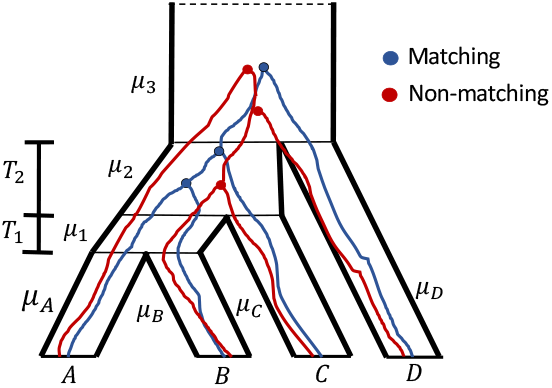
An unbalanced model species tree with four taxa with a matching and non-matching gene tree shown inside it. *T*_1_ and *T*_2_ denote the CU lengths, *µ*_*i*_s are the mutation rates, and the internal branch has an SU length of *t*_1_ = *T*_1_*µ*_1_ (the SU length for other branches are defined similarly). Taxa *A* and *B* are reffered to as cherry branches.

CASTLES uses Eq. (S5) to calculate the length of the cherry branch leading to taxa *A* (and similarly for *B*) in an unbalanced quartet species tree:

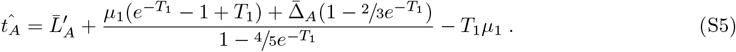

This equation depends on the mutation rate of the internal branch *µ*_1_ and the CU branch length *T*_1_. CASTLES uses the simplified formula for Δ_*I*_ in Table S5 to calculate *µ*_1_ and substitutes it in Eq. (S5):

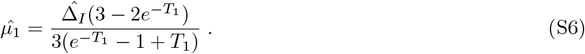

Here, 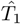, the estimate of the CU length of the internal branch, is calculated using the approach from Sayyari and Mirarab (2016). However, as *T*_1_ → 0, the denominator of Eq. (S6) becomes very small, and therefore the formula becomes unstable as 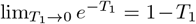. In addition, the value of *T*_1_ calculated using Sayyari and Mirarab (2016)’s algorithm degrades in accuracy as gene tree estimation error increases. While the formulas for the internal branches in CASTLES were calculated so as to not have a dependency on these CU lengths, the terminal branches still have this dependency.

To improve the estimation of *µ*_1_ and reduce the dependency on the CU branch lengths estimated using Sayyari and Mirarab (2016)’s approach in Eq. (S5), CASTLES-Pro instead first calculates *T*_1_ as a function of *δ* (see the main text) and *µ*_1_ as 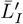. It then directly uses these values to compute the terminal branch equations.

**Table S4:**
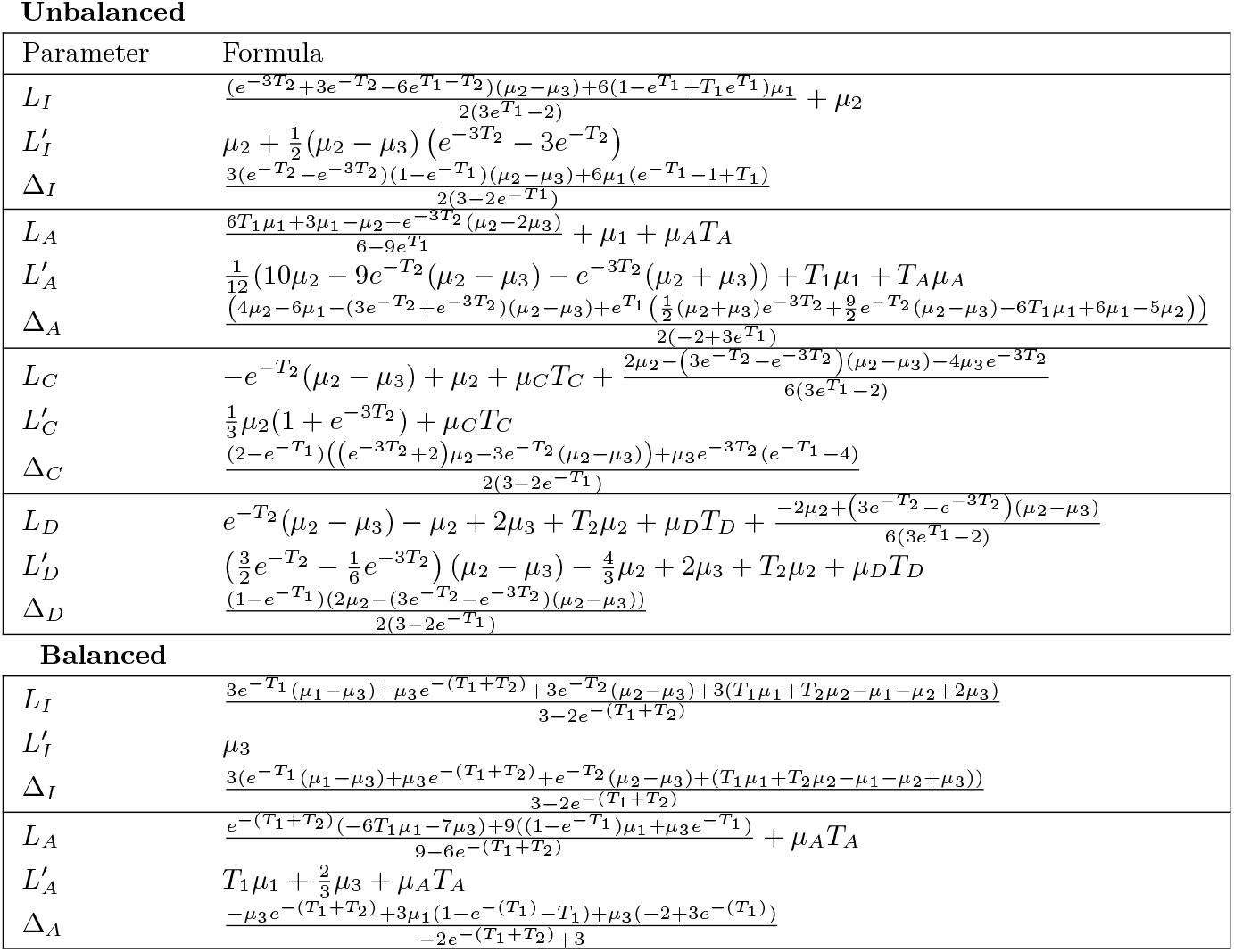
Summary of formulas for expected branch lengths in matching and non-matching quartet gene trees. This table is reproduced from Table S1 in Tabatabaee et al. (2023).

**Table S5:**
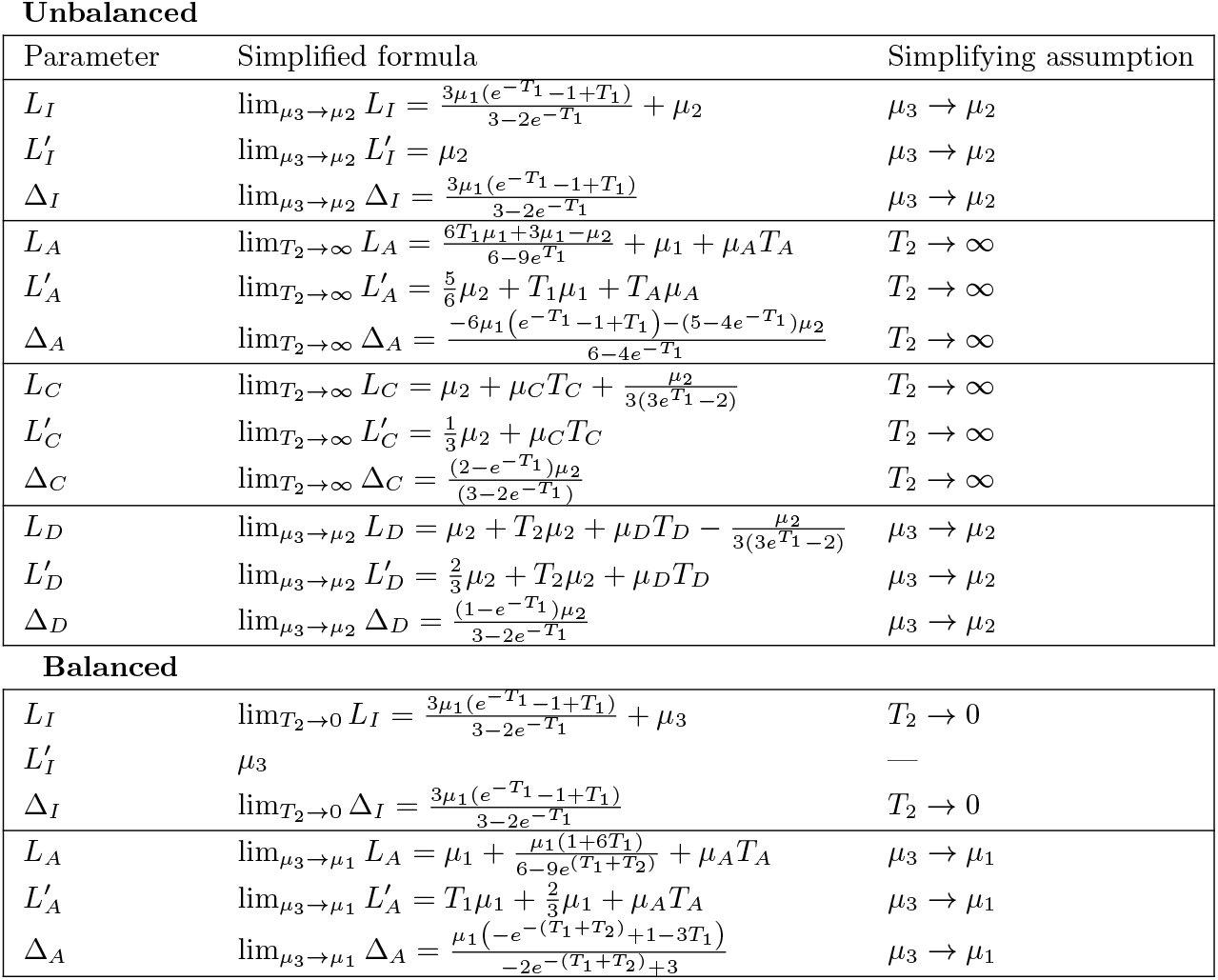
Summary of simplifying assumptions and the corresponding simplified formulas for expected branch lengths in matching and non-matching quartet gene trees. This table is reproduced from Table S2 in Tabatabaee et al. (2023).

**Table S6:** Summary of formulas for estimating unbalanced or balanced quartet species tree branch lengths in SU. This table is reproduced from Table S3 in Tabatabaee et al. (2023).

| Unbalanced |  |  |
| --- | --- | --- |
| Parameter | Estimation formula | Simplifying assumption(s) |
| $t_1$ | $\hat{t}_1 = \bar{L}'_I \left( \frac{1}{2}\bar{\delta} + \frac{1}{6}\sqrt{3\bar{\delta}(3\bar{\delta}+4)} \right)$ | $\mu_3 \rightarrow \mu_2; \mu_1 \rightarrow \mu_2$ |
| $t_A$ | $\hat{t}_A = \bar{L}'_A + \frac{\mu_1(e^{-T_1}-1+T_1)+\bar{\Delta}_A(1-2/3e^{-T_1})}{1-4/5e^{-T_1}} - T_1\mu_1$ | $T_2 \rightarrow \infty$ |
| $t_B$ | $\hat{t}_B = \bar{L}'_B + \frac{\mu_1(e^{-T_1}-1+T_1)+\bar{\Delta}_B(1-2/3e^{-T_1})}{1-4/5e^{-T_1}} - T_1\mu_1$ | $T_2 \rightarrow \infty$ |
| $t_C$ | $\hat{t}_C = \bar{L}'_C - \frac{1}{3}(2 - \frac{1}{2-e^{-T_1}})\bar{\Delta}_C$ | $T_2 \rightarrow \infty$ |
| $t_2 + t_D$ | $\hat{t}_2 + \hat{t}_D = \bar{L}'_D - \frac{2}{3}(2 + \frac{1}{1-e^{-T_1}})\bar{\Delta}_D$ | $\mu_3 \rightarrow \mu_2$ |
| Balanced |  |  |
| $t_1 + t_2$ | $\hat{t}_1 + \hat{t}_2 = \bar{L}'_I \left( \frac{1}{2}\bar{\delta} + \frac{1}{6}\sqrt{3\bar{\delta}(3\bar{\delta}+4)} \right)$ | $T_2 \rightarrow 0; \mu_1 \rightarrow \mu_3$ |
| $t_A$ | $\hat{t}_A = \bar{L}'_A - \frac{2}{3}\mu_1 - \frac{1}{3} \left( \mu_1 \left( 1 - e^{-(T_1+T_2)} \right) - \bar{\Delta}_A \left( 3 - 2e^{-(T_1+T_2)} \right) \right)$ | $\mu_3 \rightarrow \mu_1$ |
| $t_B$ | $\hat{t}_B = \bar{L}'_B - \frac{2}{3}\mu_1 - \frac{1}{3} \left( \mu_1 \left( 1 - e^{-(T_1+T_2)} \right) - \bar{\Delta}_B \left( 3 - 2e^{-(T_1+T_2)} \right) \right)$ | $\mu_3 \rightarrow \mu_1$ |
| $t_C$ | $\hat{t}_C = \bar{L}'_C - \frac{2}{3}\mu_2 - \frac{1}{3} \left( \mu_2 \left( 1 - e^{-(T_1+T_2)} \right) - \bar{\Delta}_C \left( 3 - 2e^{-(T_1+T_2)} \right) \right)$ | $\mu_3 \rightarrow \mu_2$ |
| $t_D$ | $\hat{t}_D = \bar{L}'_D - \frac{2}{3}\mu_2 - \frac{1}{3} \left( \mu_2 \left( 1 - e^{-(T_1+T_2)} \right) - \bar{\Delta}_D \left( 3 - 2e^{-(T_1+T_2)} \right) \right)$ | $\mu_3 \rightarrow \mu_2$ |

### C. Details of the Experimental Study

#### C.1 Simulated Datasets

##### ILS-only dataset

We reused a published dataset from Tabatabaee et al. (2023) simulated using Simphy. This dataset has 50 repliates, each with 100 ingroups and one outgroup species and 1000 gene trees. In addition to true gene trees, gene alignments of length 1600bp, 800bp, 400bp, and 200bp were simulated under GTR+Γ, and four sets of gene trees were estimated from these alignments using FastTree-2 (Price et al., 2010), producing gene trees with 23%, 31%, 42%, and 55% GTEE. Measuring ILS using the average Robinson and Foulds (1981) (RF) distance (AD for short) between the model species tree and true gene trees, ILS is heterogeneous across replicates and ranges from 30% to 58%, with a mean of 46% AD.

##### GDL+ILS

We updated the Simphy-generated GDL+ILS datasets of Willson et al. (2022, 2023) to have species trees with SU branch lengths. Here, locus trees evolve inside the species tree with GDL events only; the final true gene trees evolve on the locus tree under MSC. Therefore, the topological differences between the final true gene trees and the locus tree are only due to ILS; we use the normalized RF distance between the locus tree and the true gene trees to measure ILS. This dataset has model conditions (10 replicates each) characterized by two levels of ILS (low and high ILS with 20% and 65% average ILS level, respectively), six duplication rates, ranging from 10^−13^ to 10^−9^, three sequence lengths (50bp, 100bp, 500bp), four different numbers of species (21, 51, 101, or 1001), and five numbers of genes (50, 100, 500, 1000, 10,000). The loss rate varies based on the duplication rate, with three different ratios: 1 (equal loss), 0.5 or 0 (no loss). In the default model condition, the loss rate is equal to the duplication rate. Gene trees were estimated using FastTree-2. The number of replicates in all model conditions is 10. Table S7 summarizes further statistics about the model conditions of this dataset.

##### HGT+ILS

We recreated a 50-replicate 51-taxon (50 ingroup and one outgroup) dataset with both HGT and ILS based on the parameters used by Davidson et al. (2015). The ILS level is fixed at 30% AD across the model conditions. The six model conditions differ in HGT rates, leading to total discordance that varies between 30% to 68%. The average number of HGT events per gene for the six model conditions starts from 0 and increases to 0.08, 0.2, 0.8, 8 and 20 that correspond to HGT rates of 0, 2 × 10^−9^, 5 × 10^−9^, 2 × 10^−8^, 2 × 10^−7^ and 5 × 10^−7^. In addition to true gene trees, we simulated 1000bp gene sequence alignments using INDELible (Fletcher and Yang, 2009) under the GTR+Γ model and then used FastTree-2 to estimate gene trees under the GTR model. GTEE is on average 28% and about the same in all model conditions. The number of genes is 1000 and the number of replicates is 50. Table S8 summarizes the empirical statistics of this dataset.

**Table S7:** Empirical statistics of the simulated GDL datasets. AD refers to average RF distance between the locus tree and the true gene trees, and GTEE refers to average RF distance between true and estimated gene trees. L/D refers to the ratio between loss and duplication rates. The last column shows the average number of leaves in each gene family tree across the replicates. Default parameters: 1000 genes, 100bp sequence length, 10 replicates, 1 L/D ratio.

| Dup. rate | # Taxa | AD | GTEE (500bp) | GTEE (100bp) | GTEE (50bp) | L/D ratio | # Leaves |
| --- | --- | --- | --- | --- | --- | --- | --- |
| Low ILS |  |  |  |  |  |  |  |
| $1 \times 10^{-13}$ | 21 | 18.4% | – | 37.9% | – | 1 | 21.0 |
| $1 \times 10^{-12}$ | 21 | 22.1% | 18.4% | 41.5% | 52.5% | 1 | 21.0 |
| $1 \times 10^{-12}$ | 51 | 20.9% | – | 42.1% | – | 1 | 51.0 |
| $1 \times 10^{-12}$ | 101 | 25.1% | – | 45.9% | – | 1 | 101.2 |
| $1 \times 10^{-11}$ | 21 | 25.1% | – | 42.2% | – | 1 | 21.3 |
| $1 \times 10^{-10}$ | 21 | 19.0% | – | 36.8% | – | 1 | 24.1 |
| $1 \times 10^{-10}$ | 101 | 23.4% | – | 44.0% | – | 1 | 116.6 |
| $1 \times 10^{-10}$ | 101 | 23.5% | – | 45.0% | – | 0.5 | 128.0 |
| $1 \times 10^{-10}$ | 101 | 24.0% | – | 44.1% | – | 0 | 145.1 |
| $5 \times 10^{-10}$ | 21 | 15.8% | – | 34.2% | – | 1 | 35.8 |
| $5 \times 10^{-10}$ | 101 | 20.3% | 19.29% | 43.36% | 55.68% | 1 | 165.3 |
| $5 \times 10^{-10}$ | 101 | 22.71% | – | 45.08% | – | 0.5 | 290.6 |
| $5 \times 10^{-10}$ | 101 | 26.59% | – | 43.09% | – | 0 | 550.0 |
| $5 \times 10^{-10}$ | 1001 | 23.49% | – | 44.4% | – | 1 | 1578.1 |
| $1 \times 10^{-9}$ | 21 | 15.6% | – | 33.7% | – | 1 | 52.1 |
| $1 \times 10^{-9}$ | 101 | 19.1% | – | 39.7% | – | 1 | 228.5 |
| $1 \times 10^{-9}$ | 101 | 20.2% | – | 43.4% | – | 0.5 | 993.0 |
| $1 \times 10^{-9}$ | 101 | 23.9% | – | 47.2% | – | 0 | 3727.8 |
| High ILS |  |  |  |  |  |  |  |
| $1 \times 10^{-13}$ | 21 | 69.3% | – | 42.2% | – | 1 | 21.0 |
| $1 \times 10^{-12}$ | 21 | 67.4% | 19.2% | 42.6% | 55.7% | 1 | 21.0 |
| $1 \times 10^{-12}$ | 51 | 75.7% | – | 45.8% | – | 1 | 51.2 |
| $1 \times 10^{-12}$ | 101 | 78.4% | – | 48.2% | – | 1 | 101.2 |
| $1 \times 10^{-11}$ | 21 | 67.0% | – | 41.0% | – | 1 | 21.3 |
| $1 \times 10^{-10}$ | 21 | 64.5% | – | 39.0% | – | 1 | 24.2 |
| $5 \times 10^{-10}$ | 21 | 54.5% | – | 39.5% | – | 1 | 36.2 |
| $5 \times 10^{-10}$ | 101 | 50.0% | – | 43.9% | – | 1 | 170.1 |
| $1 \times 10^{-9}$ | 21 | 44.2% | – | 38.3% | – | 1 | 47.9 |

**Table S8:** Empirical statistics of the simulated HGT datasets. AD refers to average RF distance between the model species tree and true gene trees, and GTEE refers to average RF distance between true and estimated gene trees. The number of taxa is 51 and the number of genes is 1000.

| HGT rate | Expected number of HGT events per gene | AD | GTEE |
| --- | --- | --- | --- |
| 0 | 0 | 29.7% | 27.0% |
| $2 \times 10^{-9}$ | 0.08 | 30.1% | 31.2% |
| $5 \times 10^{-9}$ | 0.2 | 30.9% | 28.9% |
| $2 \times 10^{-8}$ | 0.8 | 34.1% | 28.4% |
| $2 \times 10^{-7}$ | 8 | 53.4% | 27.0% |
| $5 \times 10^{-7}$ | 20 | 68.4% | 26.9% |

### C.2 Biological Datasets

#### Birds

We studied the birds dataset of Stiller et al. (2024) including 363 species and 63,430 genes that was used to resolve family-level relationships among neoavian species and is expected to have high levels of ILS due to a rapid radiation. The original study had inferred the tree topology using ASTRAL and then estimated branch lenghts on that topology using the concatenation of all 63K genes. We infer branch lengths on the same ASTRAL topology using CASTLES-Pro.

#### Bees

We renalyzed the bees dataset of Bossert et al. (2021) containing 32 species (30 ingroups and two outgroups) from the bee subfamily Nomiinae and 853 gene trees (estimated using RAxML). We used the ASTRAL topology from the original study, and used partitioned concatenation (with RAxML) and CASTLES-Pro to draw branch lengths on this topology.

#### Mammals

We studied the mammalian biological dataset from Song et al. (2012), including 37 species (36 ingroup and 1 outgroup) and 447 gene trees, which was reduced to 424 trees after removing gene trees with mis-matching names (Mirarab et al., 2014). We estimated an ASTRAL topology and estimated branch lengths using concatenation and CASTLES-Pro on that topology.

#### Fungi

We examined the fungal dataset of Butler et al. (2009), including 16 yeast species and 7,180 multi-copy gene family trees. The original study had used MrBayes (Huelsenbeck and Ronquist, 2001) on a concatenated alignment created by sampling 30,000 sites from 706 individual gene family orthologous peptide sequences. We used ASTRAL-Pro2 (Zhang and Mirarab, 2022a) to estimate a species tree using all 7,180 gene family trees, and used CASTLES-Pro to estimate branch lengths on that tree. The two trees are different in one branch, with an RF distance of 7.6%.

#### Plants (1KP)

We analyzed the plants dataset of Wickett et al. (2014), that included 103 species and 424 single-copy gene trees, as well as 9,610 multi-copy gene family trees for 83 of the species, that was left unsued in the original study due to lack of propor method for estimating the species tree from multi-copy input. The gene trees were inferred using RAxML for the first two codon positions (C12) in the transcriptome. We compare the branch lengths of the concatenation tree inferred from the 424 single-copy gene alignments with a tree inferred using ASTRAL-Pro2 from all 9,610 multi-copy gene trees furnished with CASTLES-Pro branch lengths. The two trees have 80 taxa in common and are different in 7 branches on the shared set of taxa, resulting in an RF distance of 9.1%.

#### Eudicots

We studied the 40-taxon angiosperm dataset of Chanderbali et al. (2022) focused on the Eudicots lineage. This study had used three sets of genes to perform phylogenomic analysis using concatenation and coalescent-based summary methods: 345 filetered single-copy Angiosperms353 loci (Johnson et al., 2019), 1248 single-copy BUSCO (Simão et al., 2015) genes, and 2,573 multi-copy orthogroups. The authors had performed concatenation analysis on the two sets of single-copy genes (Angiosperms353 loci and BUSCOs) using RAxML and coalescent analysis on all three sets of genes using ASTRAL and ASTRAL-Pro for single and multi-copy input respectively. We compared the two concatenation trees from the original study with a tree we inferred using ASTRAL-Pro2 from the 2,573 orthogroups that was furnished with CASTLES-Pro branch lengths. The two concatenation trees had the same topology that was different from the ASTRAL-Pro2 tree in three branches, with an RF distance of 8.1%.

#### Microbial datasets

We analyzed three microbial datasets including thousands of species of bacteria and archaea and different sets of genes to study a debate about the length of the branch separating domains archaea and bacteria (AB branch). While the long-standing hypothesis was that these two domains are separated by a long branch (Cox et al., 2008; Gogarten et al., 1989; Iwabe et al., 1989), a recent study (Zhu et al., 2019) had estimated a far shorter length for the AB branch than what was previously expected using a concatenation analysis. Moody et al. (2022a) had further studied this and other bacterial datasets, and suggested that concatenation can severaly underestimate branch lengths on datasets with high levels of HGT, resulting in short estimates of Zhu et al. (2019).

Here we examine two bacterial datasets analyzed by Moody et al. (2022a) and the Web of Life (WoL) dataset from Zhu et al. (2019) with CASTLES-Pro to further study this debate. The two bacterial datasets include a 72-taxon dataset with 49 core genes originally from Williams et al. (2020) that includes ribosomal proteins and other conserved elements and a 108-taxon dataset with 38 genes from Petitjean et al. (2015) that only includes non-ribosomal proteins. The WoL dataset included 10,575 species (9,906 bacteria and 669 archaea) and 381 marker genes including ribosomal and non-ribosomal proteins. For the two smaller bacterial datasets, we estimated a species tree using ASTRAL on the two gene sets and used concatenation (with RAxML) and CASTLES-Pro to draw branch lengths on these topologies. On the WoL dataset, we used the ASTRAL tree from the original study that was furnished with branch lengths estimated using RAxML from a concatenation including 100 sites randomly selected from sites with less than 50% gaps for each marker gene. We estimated branch lengths on the same topology using CASTLES-Pro.

**Table S9:** Empirical statistics of the biological datasets. ILS, GDL and HGT refer to the main source of gene tree discordance in each dataset.

| Study | Dataset type/species | # taxa | # single-copy genes | # multi-copy genes |
| --- | --- | --- | --- | --- |
| **ILS** |  |  |  |  |
| Stiller et al. (2024) | Neoavian birds | 363 | 63,430 | NA |
| Song et al. (2012) | Mammals | 37 | 424 | NA |
| Bossert et al. (2021) | Bees (subfamily Nomiinae) | 32 | 853 | NA |
| **GDL** |  |  |  |  |
| Wickett et al. (2014) | Plants (1kp) | 80 | 424 | 9,610 |
| Chanderbali et al. (2022) | Eudicots | 40 | 345 | 2,573 |
| Butler et al. (2009) | Fungi | 16 | 706 | 7,180 |
| **HGT** |  |  |  |  |
| Williams et al. (2020) | Bacterial (core genes) | 72 | 49 | NA |
| Petitjean et al. (2015) | Bacterial (non-ribosomal genes) | 108 | 38 | NA |
| Zhu et al. (2019) | Bacterial (WoL) | 10,575 | 381 | NA |

### C.3 Methods and Software Commands

Here we bring the details of the methods and software commands. All experiments were performed on the University of Illinois campus cluster, with a memory limit of 128GB.

#### DISCO

We used DISCO (Willson et al., 2022) version 1.3 to decompose multi-copy gene-family trees into single-copy gene trees. DISCO is available at https://github.com/JSdoubleL/DISCO. We used the following command:

~~~
python 3 disco . py -i <multi - copy - gene - trees > -o <single - copy - gene - trees > -d _
~~~

#### CA-DISCO

To run concatenation on multi-copy gene family sequences, we used the script ca disco.py available at https://github.com/JSdoubleL/DISCO with the following command:

~~~
ca_disco . py -i < gene_tree_path > -a < alignment_list_path > -t < taxa_list_path >
-o <output_path > -d _
~~~

where <gene tree path> is the path to the set of multi-copy gene family trees, <alignment list path> is a file containing the list of individual sequence alignments for each gene family, and <taxa list path> is the set of all taxa. The output is the concatenated sequence alignment, that is then passed to RAxML (v8.2.12) (Stamatakis, 2014), available at https://github.com/stamatak/standard-RAxML, to optimize branch lengths on a fixed tree topology with the option -f e using the following command.

~~~
raxmlHPC - PTHREADS -f e -t < species_tree_path > -m GTRGAMMA -s < alignment_path >
-n RES -p 4321 -T 16
~~~

#### ERaBLE

To run ERaBLE (Binet et al., 2016), we first calculated a matrix of pairwise patristic distances per gene using a custom script available at https://github.com/ytabatabaee/CASTLES-Pro-paper/scripts/patristic_dist_matrix.py that uses calculate.treecompare._get_length_diffs from the package DendroPy (Sukumaran and Holder, 2010). When calculating the patristic distance matrix, we impute missing values with averages: i.e., if two taxa *i* and *j* do not appear in the same gene tree *g* together due to missing taxa in genes, as is the case in DISCO gene trees, the patristic distance of *i* and *j* in the matrix for gene tree *g* is replaced by the average patristic distances of these two taxa in the rest of the gene trees where they appear together. We used the following command to run this script

~~~
python 3 patristic_dist_matrix . py -g < gene_tree_path > -o < dist_mat. phylip >
-m all
~~~

Then we ran ERaBLE (v1.0) available at http://www.atgc-montpellier.fr/erable/ with the following command;

~~~
erable -i < dist_mat. phylip > -t < species_tree_path > -o <output_path >
~~~

#### FastME

Similar to ERaBLE, to run FastME (Lefort et al., 2015), we first computed a *single* distance matrix corresponding to average patristic distances between pairs of taxa using the following command

~~~
python 3 patristic_dist_matrix . py -g < gene_tree_path > -o < dist_mat. phylip >
-m avg
~~~

and then we ran FastME version 2.1.6.2 with the following command

~~~
fastme -2.1.6.2 - linux64 -i < dist_mat. phylip > -w BalLS -u < species_tree_path >
-o <output_path >
~~~

#### CASTLES-Pro

CASTLES-Pro is integrated inside the species tree estimation software ASTER that is available at https://github.com/chaoszhang/ASTER. To infer branch lengths on a fixed tree topology using ASTER (v1.19.3.5), we used the following commands for multi-copy and single-copy gene trees respectively

~~~
bin / astral - pro2 -i <gene - tree - path > -C -c <species - tree - topology > -o <output - path >
-- root <outgroup - name > -- genelength <gene - sequence - length >
bin / astral4 -i <gene - tree - path > -C -c <species - tree - topology > -o <output - path >
-- root <outgroup - name > -- genelength <gene - sequence - length >
~~~

where --root specifies the outgroup name (if known) and --genelength specifies the average gene sequence length (default: 1000bp).

#### CASTLES

CASTLES (Tabatabaee et al., 2023) is available at https://github.com/ytabatabaee/CASTLES. To run it, we first annotated the species tree toplogy using ASTER with the following command

~~~
astral -C -i <gene - tree - path > -c <species - tree - topology > -o <output_path >
-- root <outgroup - name > > < annotated . tre >
~~~

where the annotated tree is printed to the file <annotated.tre>. We then ran the CASTLES script (v1.0) using the following command

~~~
python 3 castles. py -t < annotated . tre > -g < gene_tree_path > -o <output_path >
~~~

#### TCMM

TCMM (Arasti et al., 2024) is available at https://github.com/shayesteh99/TCMM. To run the per-gene version, we used the command:

~~~
python 3 multiple_tree_matching . py -i <species - tree - topology > -r < gene_tree_path >
-l <lambda > -o <output_path >
~~~

To run the consensus version of TCMM, we used the command:

~~~
python 3 weighted_tree_matching . py -i <species - tree - topology > -r < gene_tree_path >
-l <lambda > -o <output_path >
~~~

### D. Additional Results (Figures and Tables)

**Figure S4:**
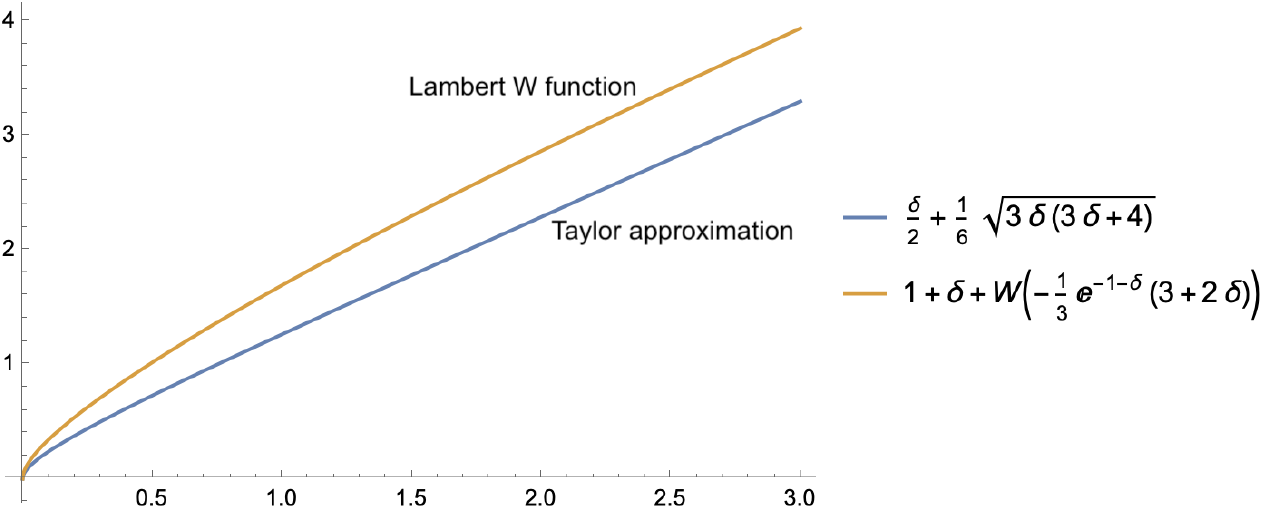
Lambert W function vs its Taylor approximation for caluclating the length of the internal branch 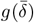 in CASTLES-Pro for *δ* ∈ (0, 3).

**Figure S5:**
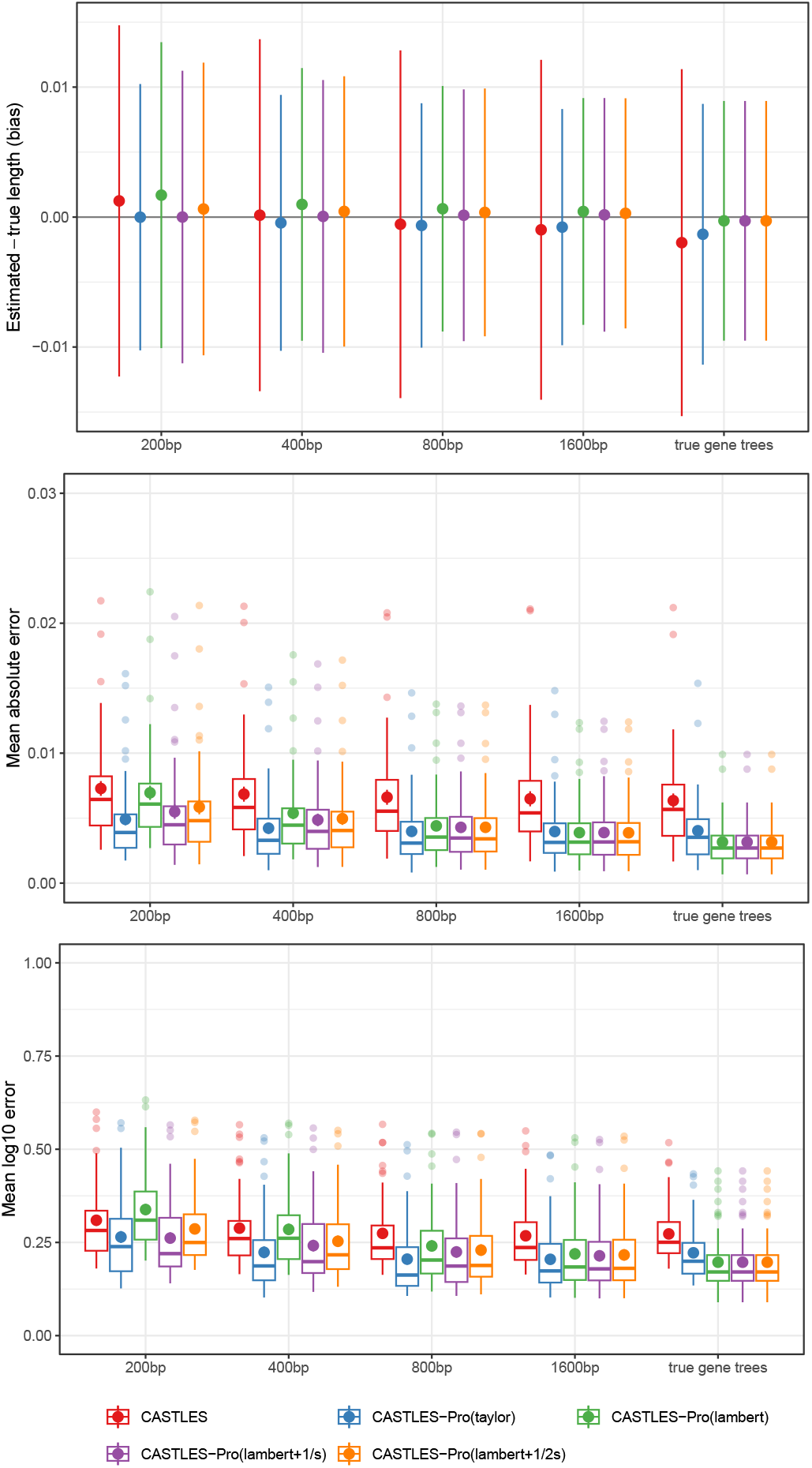
Bias, mean absolute error and mean log error for variants of CASTLES-Pro and CASTLES on 100-taxon simulated ILS datasets. The four variants of CASTLES-Pro either use the taylor approximation for calculating the length of the internal branch, or lambert function with two different pseudo-counts based on sequence lengths. The average ILS level on this dataset is 47% AD and the GTEE level varies between 0% for true gene trees to 55% for gene trees estimated from 200bp alignments. The number of genes is 1000 and the number of replicates is 50. The method shown in purple (CASTLES-Pro(lambert+1/s)) is the variant that we selected. The difference between CASTLES and CASTLES–Pro(taylor) is the way they compute mutation rates (see Supplementary Section B).

**Figure S6:**
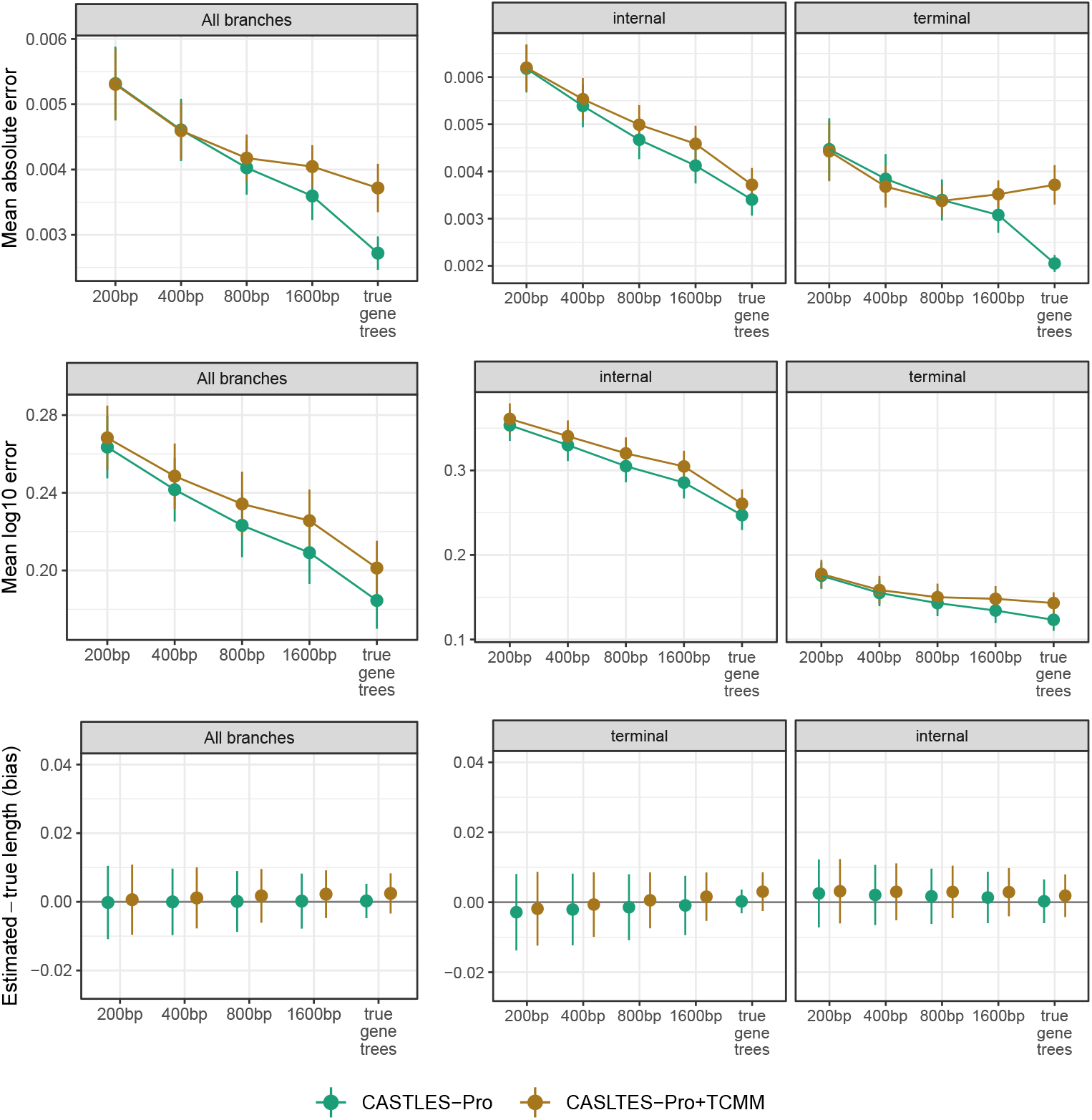
Bias, mean absolute error and mean log error for CASTLES-Pro and CASTLES-Pro+TCMM on 100-taxon simulated ILS datasets. The average ILS level on this dataset is 47% AD and the GTEE level varies between 0% for true gene trees to 55% for gene trees estimated from 200bp alignments. The number of genes is 1000 and the number of replicates is 50.

**Figure S7:**
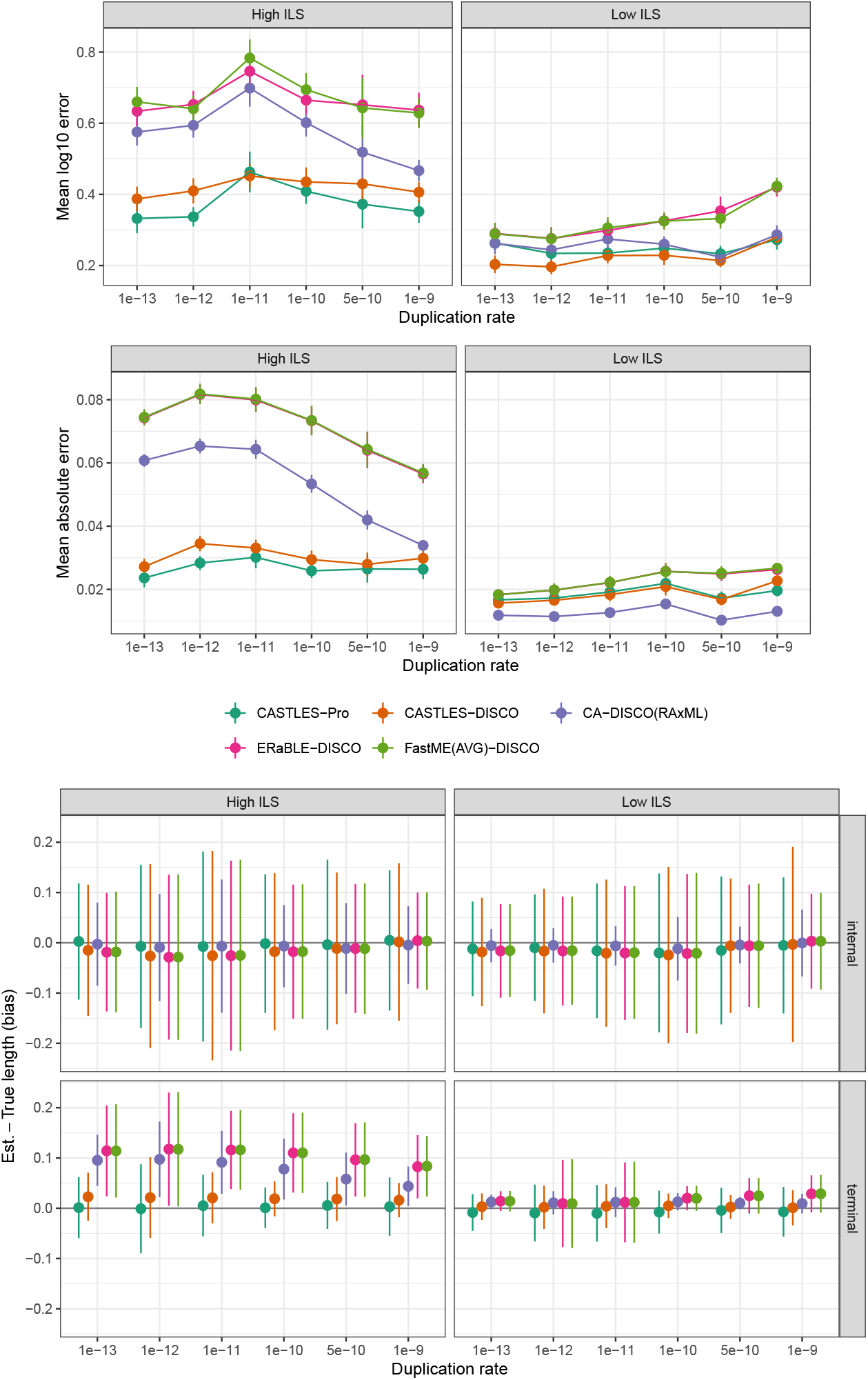
Mean log error, mean absolute error and bias on the GDL datasets for varying duplication rates. The gene trees are estiamted from 100bp alignments with average GTEE levels that varies between 33.7% to 42.2% for the low ILS condition and 38.3% to 42.6% for the high ILS condition. The number of taxa is 20, the number of genes is 1000 and the number of replicates is 10.

**Figure S8:**
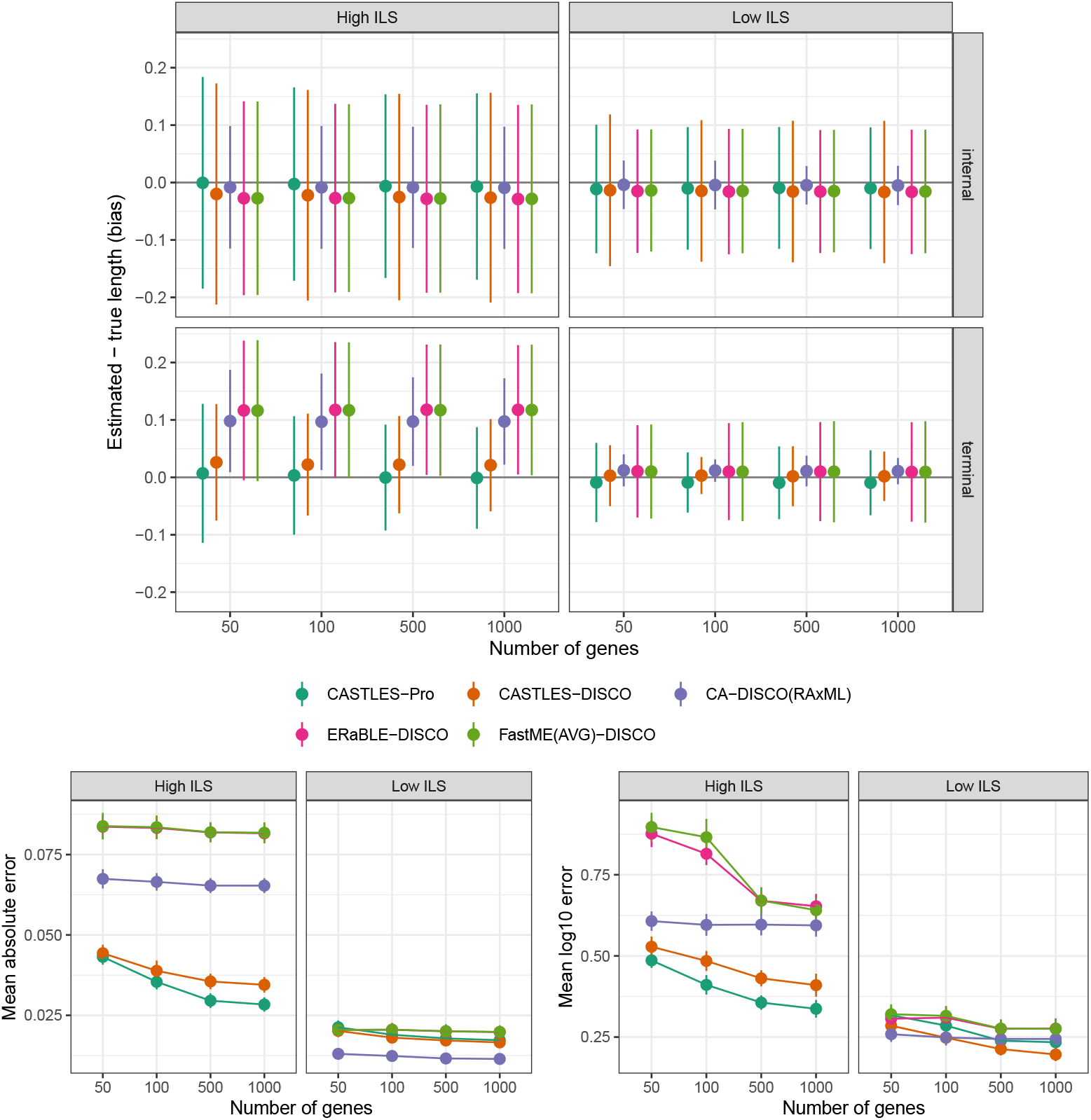
Bias, mean absolute error and mean log error for simulated GDL datasets for varying number of genes. The duplication rate is 10^−12^ with an equal loss rate. Gene trees are estiamted from 100bp alignments and the average GTEE rates for 1000 genes for the low ILS and high ILS conditions are 41.5% and 42.6% respectively. The number of taxa is 20 and the number of replicates is 10.

**Figure S9:**
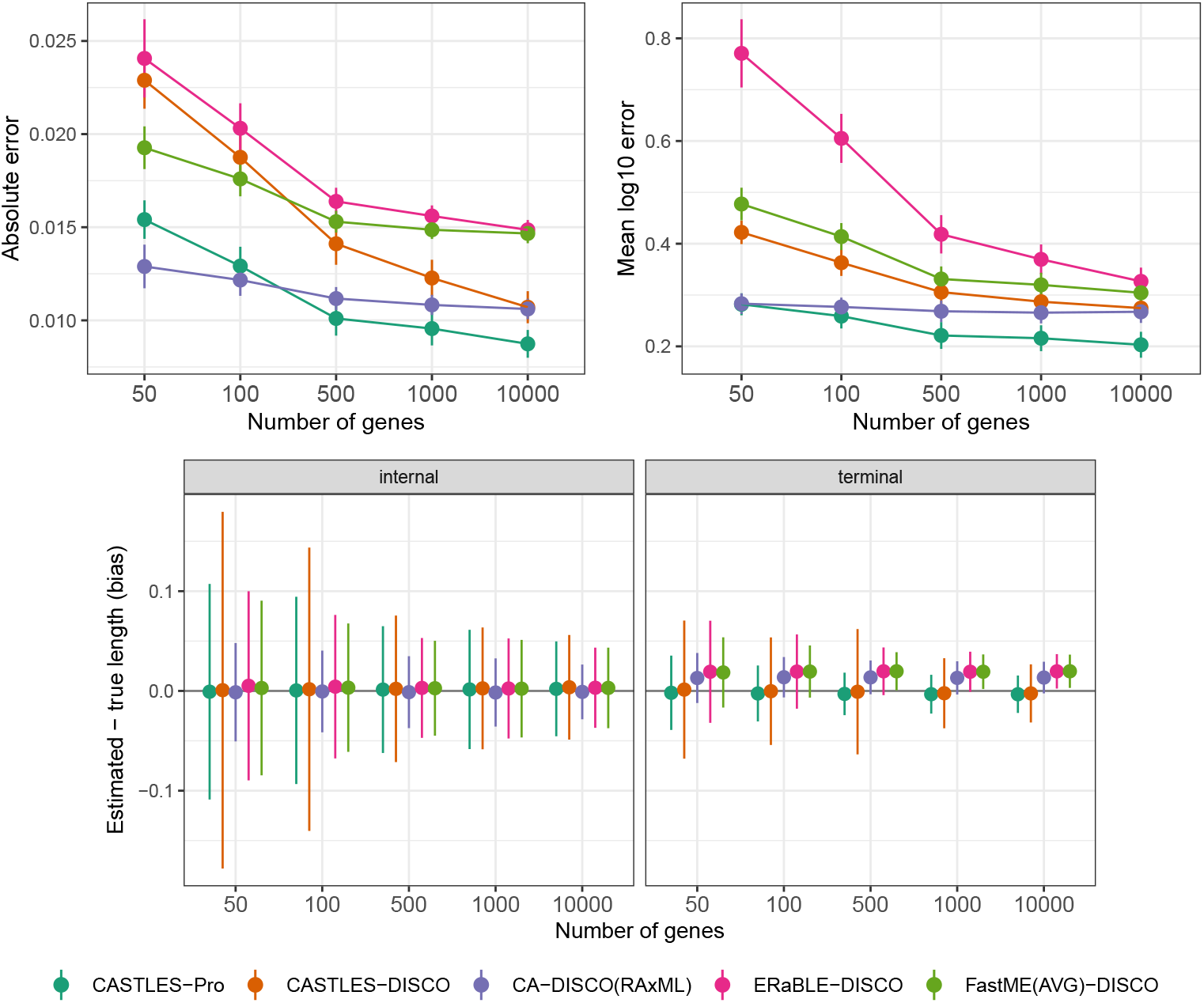
Mean log error, mean absolute error and bias on the 100-taxon GDL datasets for different number of genes. The duplication rate is 5 *×* 10^−10^ with equal loss rate and the level of ILS is low (20.3% AD). Gene trees are estiamted from 100bp alignments with an average GTEE level of 41.1% for 10,000 genes. The number of replicates is 10.

**Figure S10:**
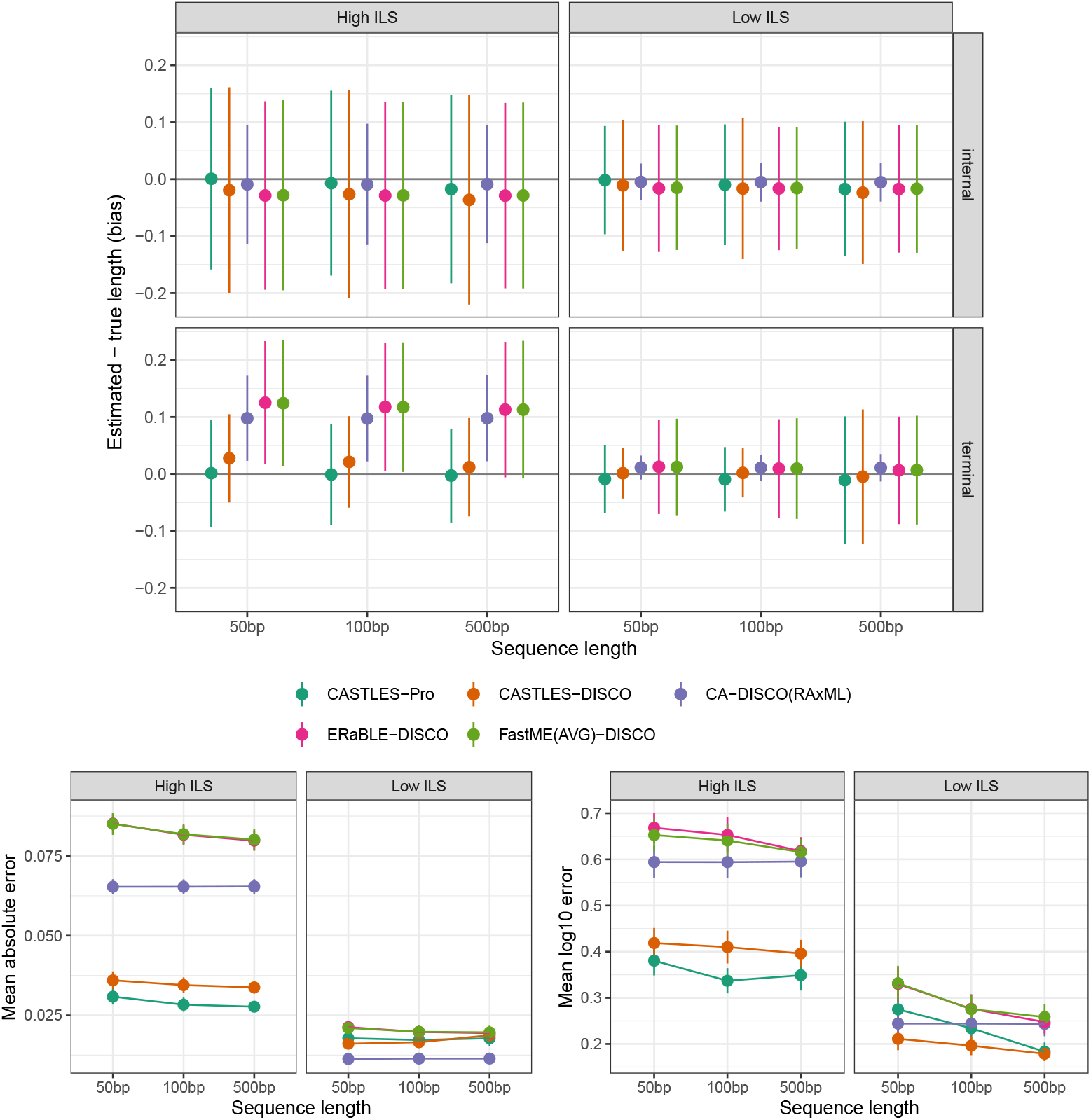
Bias, mean absolute error and mean log error for simulated GDL datasets for varying sequence lengths. The duplication rate is 10^−12^ with an equal loss rate. The average GTEE rates for the 50bp, 100bp and 500bp alignments for the low ILS condition are 52.5%, 41.5% and 18.4% respectively and for the high ILS condition are 55.7%, 42.6% and 19.2%. The number of taxa is 20, the number of genes is 1000 and the number of replicates is 10.

**Figure S11:**
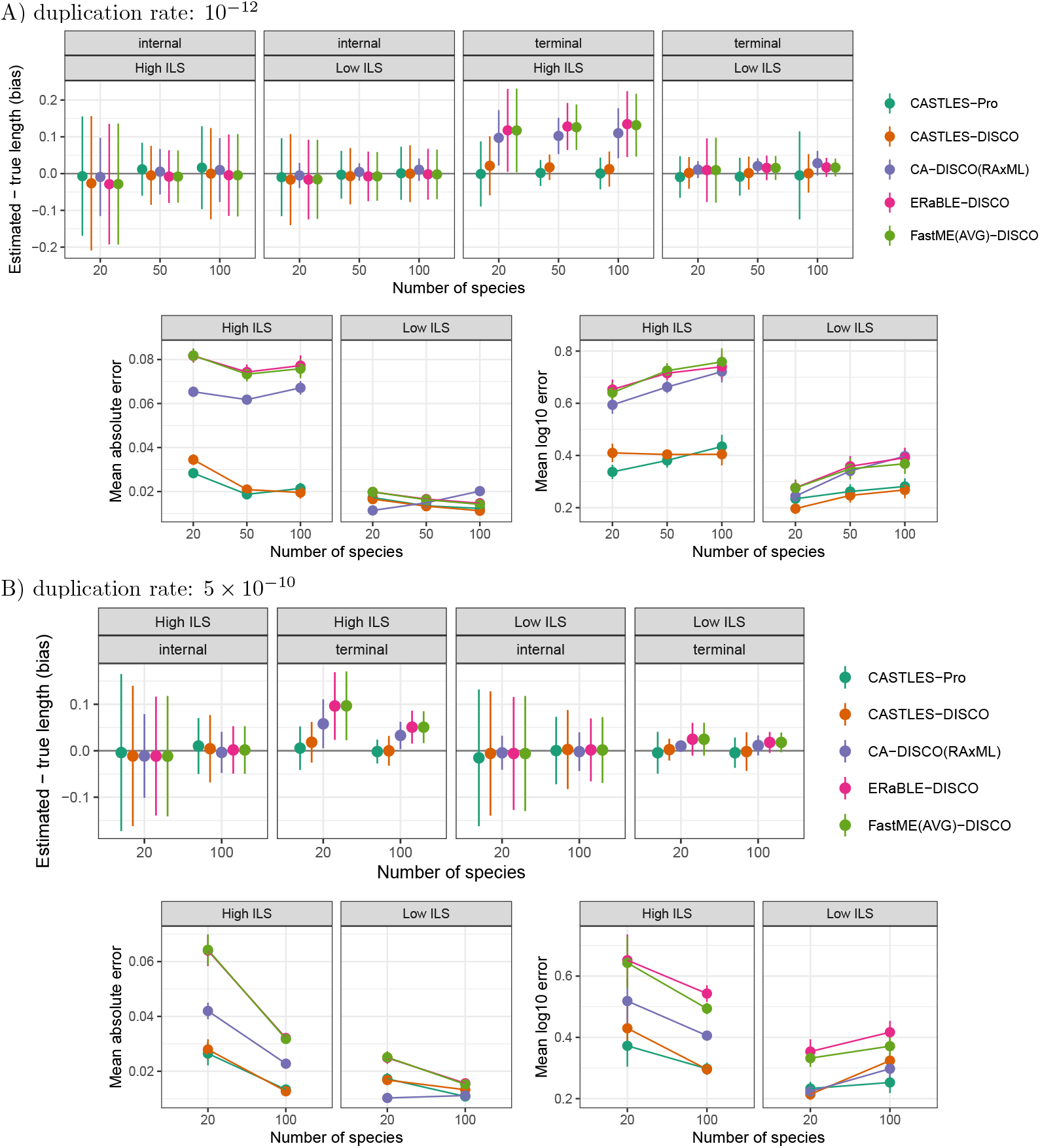
Bias, mean absolute error, and mean log error for simulated GDL datasets for varying numbers of species and level of ILS. The duplication rate is 10^−12^ (A) or 5 *×* 10^−10^ (B) with an equal loss rate. The gene trees are estimated from 100bp alignments with an average GTEE level that varies between 34.2% to 48.2% for different conditions. The number of genes is 1000, and the number of replicates is 10.

**Figure S12:**
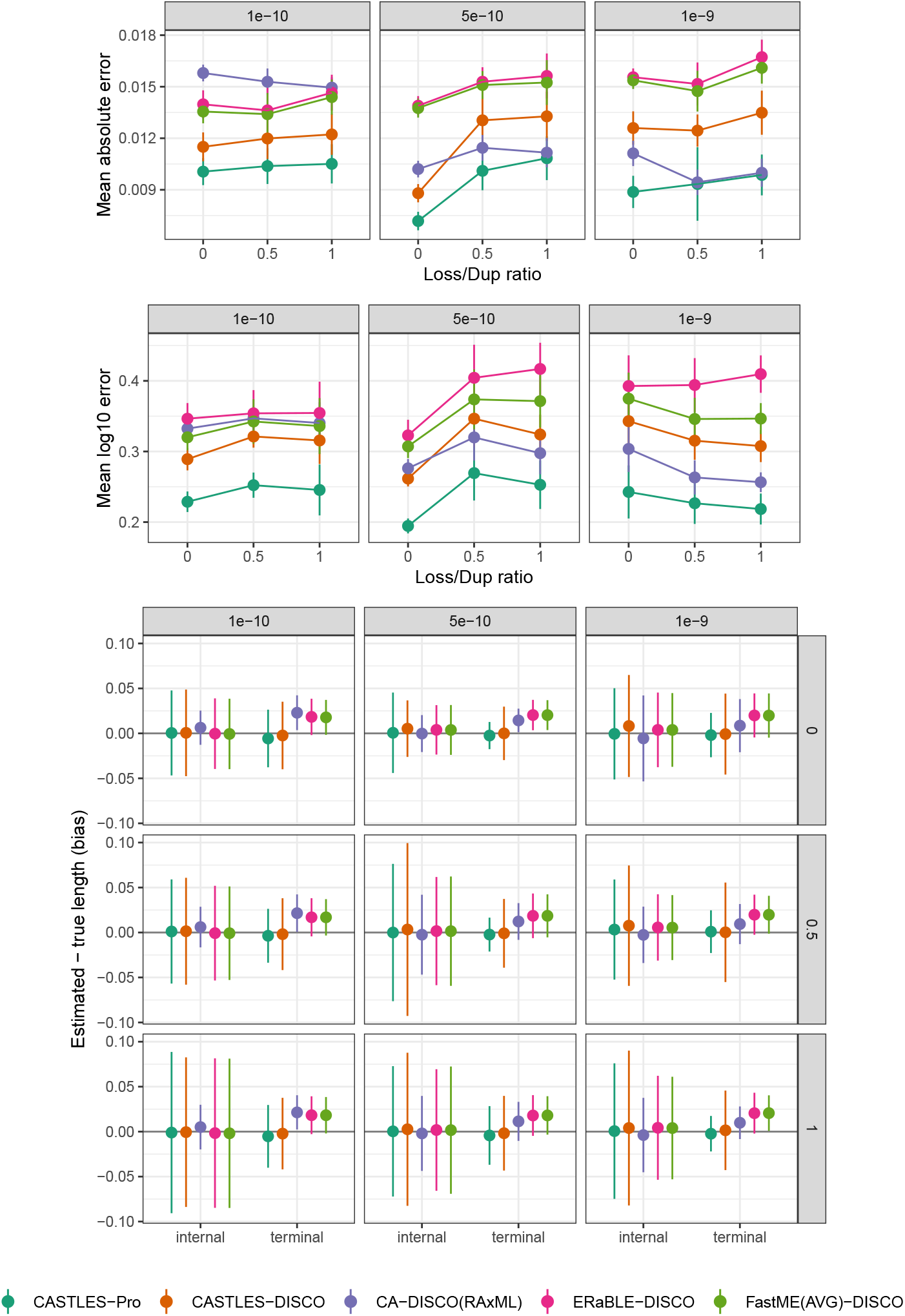
Mean log error, mean absolute error and bias on the 100-taxon GDL datasets for varying duplication/loss ratios. The gene trees are estiamted from 100bp alignments with average GTEE levels that varies between 39.7% to 47.2%. The level of ILS varies between 19.1% to 26.6%. The number of taxa is 100, the number of genes is 1000 and the number of replicates is 10.

**Figure S13:**
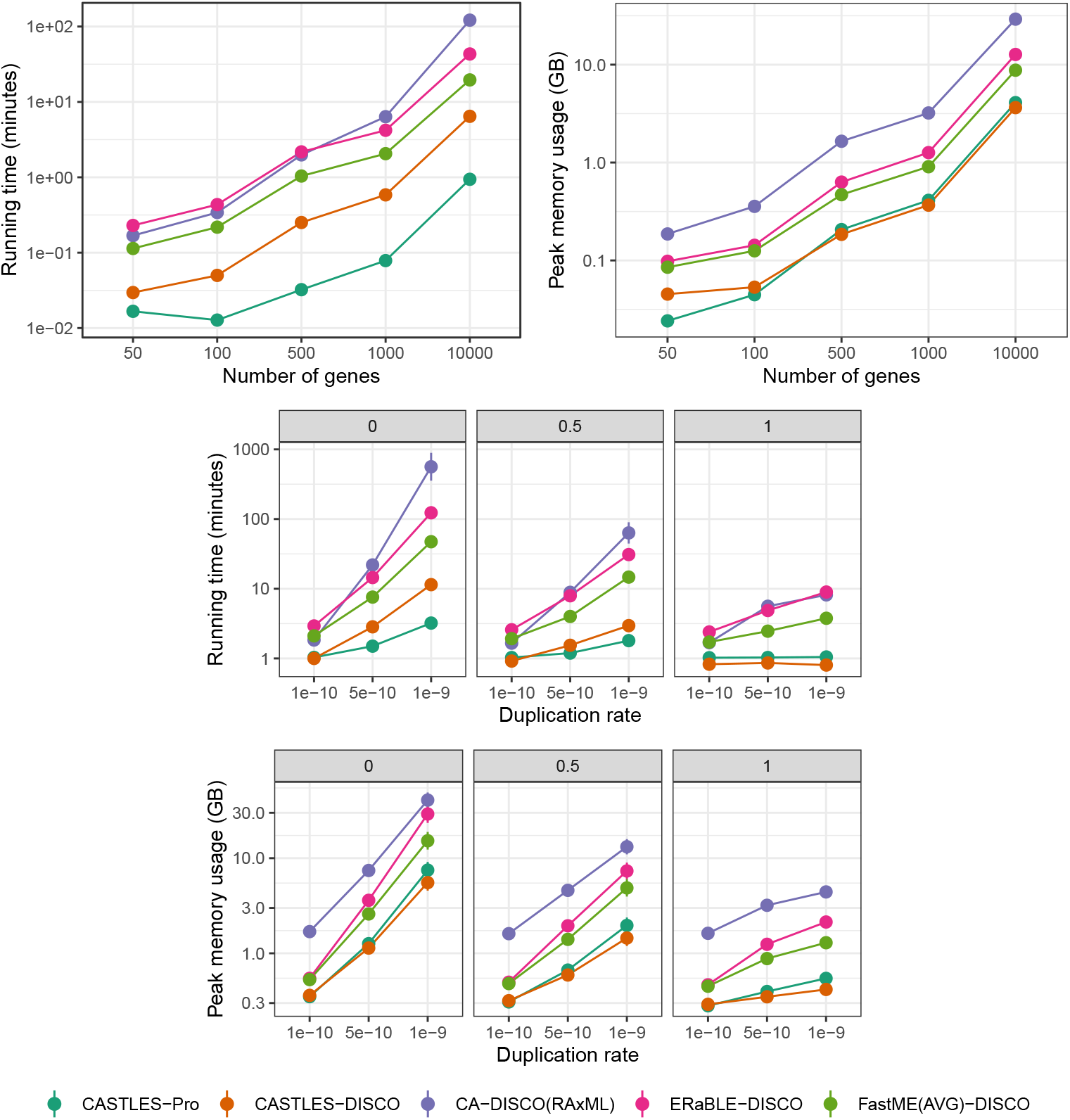
Runtime and peak memory usage of branch length estimation methods on 100-taxon GDL datasets for different number of genes and duplication rates. Gene trees are estimated from 100bp sequence alignments. The y-axes are shown in log scale. (top) The duplication rate is 5 *×* 10^−10^ with equal loss rate and the number of genes varies between 50 to 10,000. (bottom) The duplication rate varies between 10^−10^ to 10^−9^ and the number of genes is 1000. The panels show L/D ratio. The number of replicates is 10. The runtime does not include gene tree estimation or species tree topology estimation time, as all methods draw branch lengths on a fixed species tree topology.

**Figure S14:**
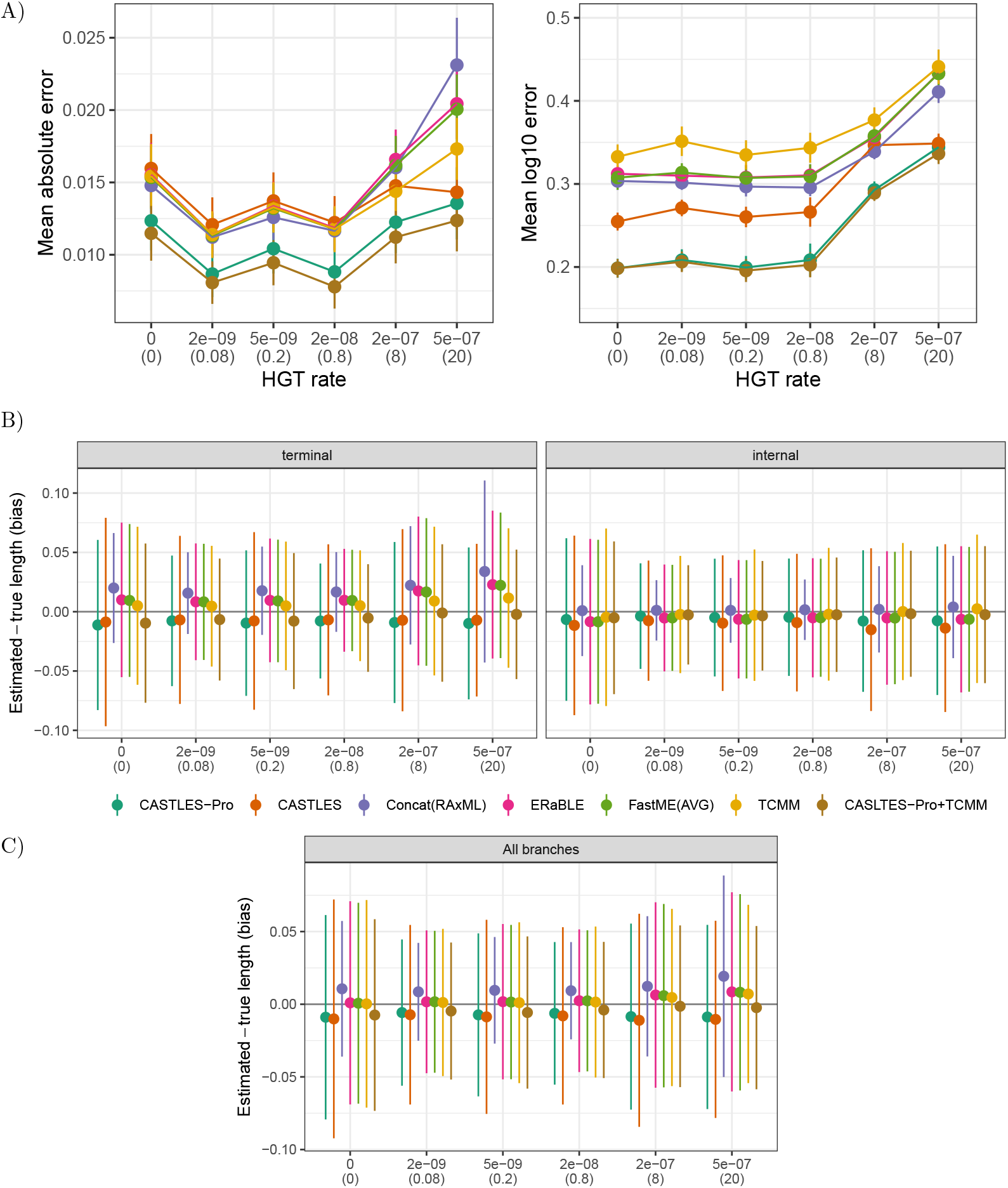
Mean absolute error, mean log error, and bias of branch length estimation methods on 50-taxon simulated HGT datasets. The x-axis indicates the rate of HGT and the expected number of HGT events per gene (in parentheses). The number of replicates is 50, and the number of genes is 1000.

**Figure S15:**
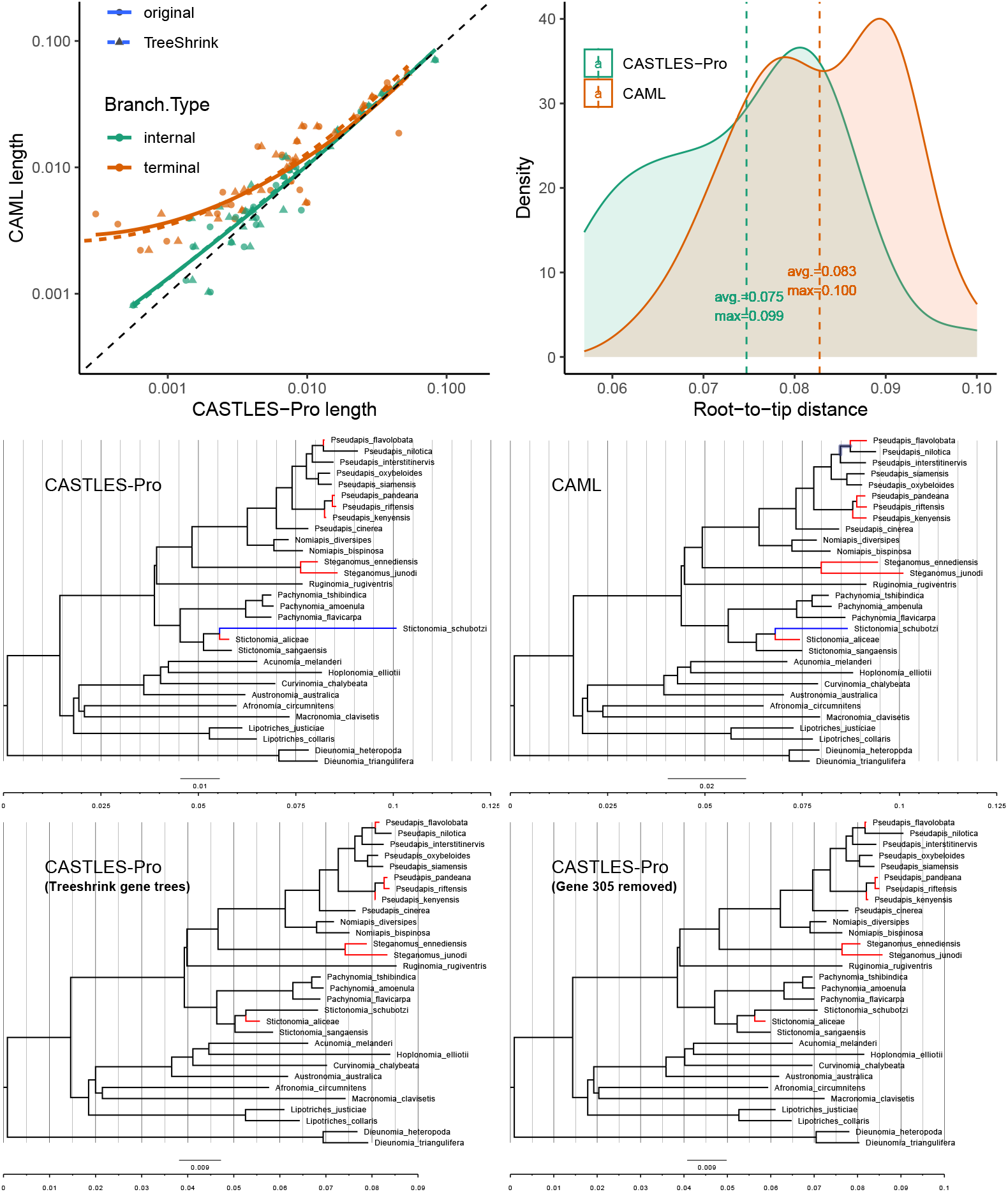
Comparison between the branch lengths of CASTLES-Pro (with or without TreeShrink) and CAML on the bees dataset of Bossert et al. (2021) after removing the outgroup taxa *Lasioglossum albipes* and *Dufourea novaeanglia*. The number of species (after removing the outgroups) is 30 and the number of gene trees is 853. We used the ASTRAL topology from the original study, and used concatenation and CASTLES-Pro to draw branch lengths on this topology. We set the average gene sequence length in CASTLES-Pro as 650bp, as the concatenated alignment had 867 loci with 576,041bp length in total, with an average loci length of 664.4. The branch lengths that are at least 2x longer or shorter in CASTLES-Pro tree compared to the concatenation tree are highlighted in blue and red respectively. The CASTLES-Pro tree estimates an unusually long branch length for taxon *Stictonomia schubotzi* due to an outlier gene, that is fixed when using TreeShrink gene trees or gene trees with the outlier gene 305 removed.

**Figure S16:**
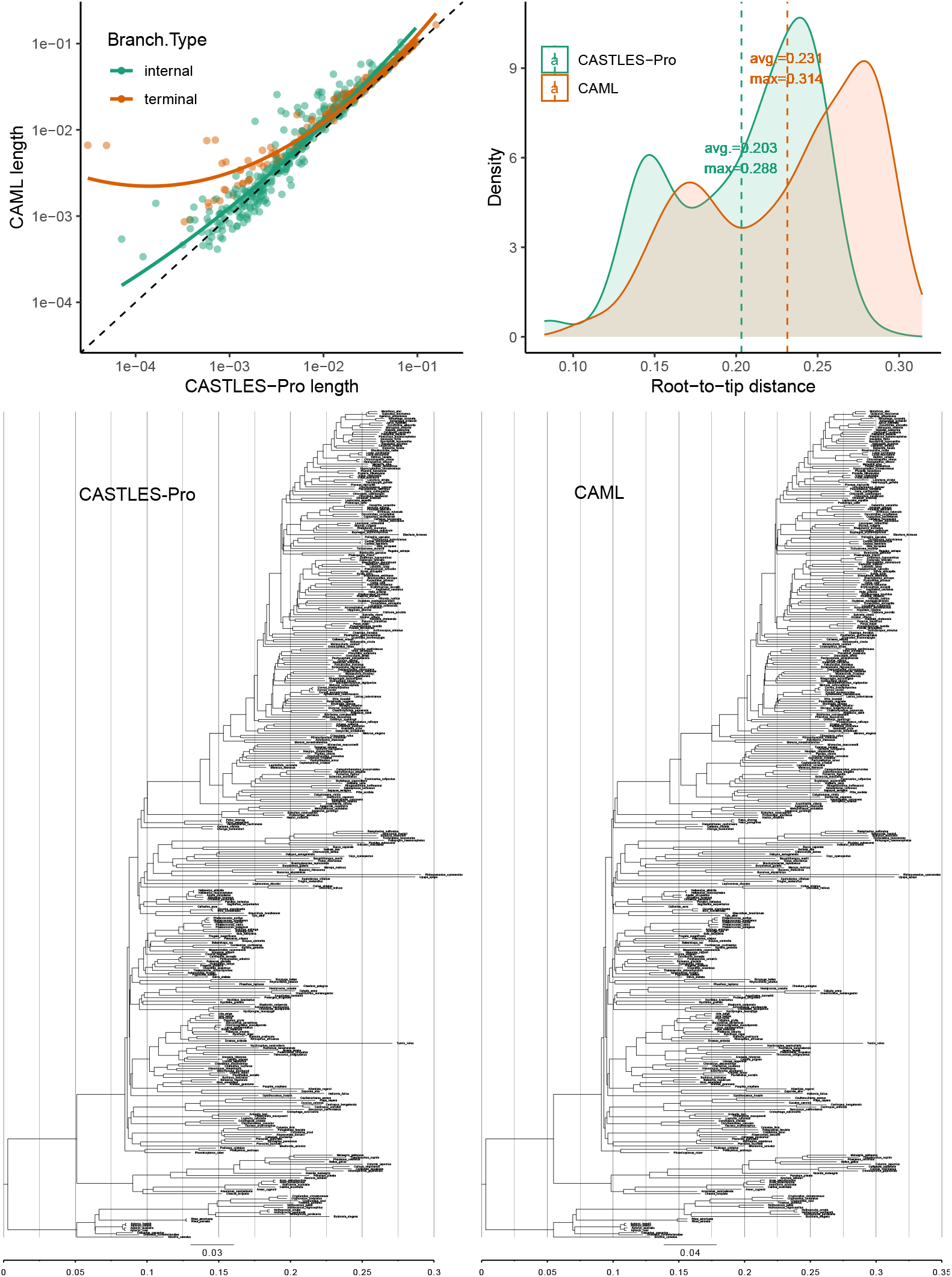
Comparison between branch lengths of CASTLES-Pro and CAML on the birds datasets of Stiller et al. (2024). The number of species is 363 and the number of gene trees is 63,430. We used the ASTRAL topology from the original study with concatenation branch lengths, and estimated branch lengths with CASTLES-Pro on the same topology.

**Figure S17:**
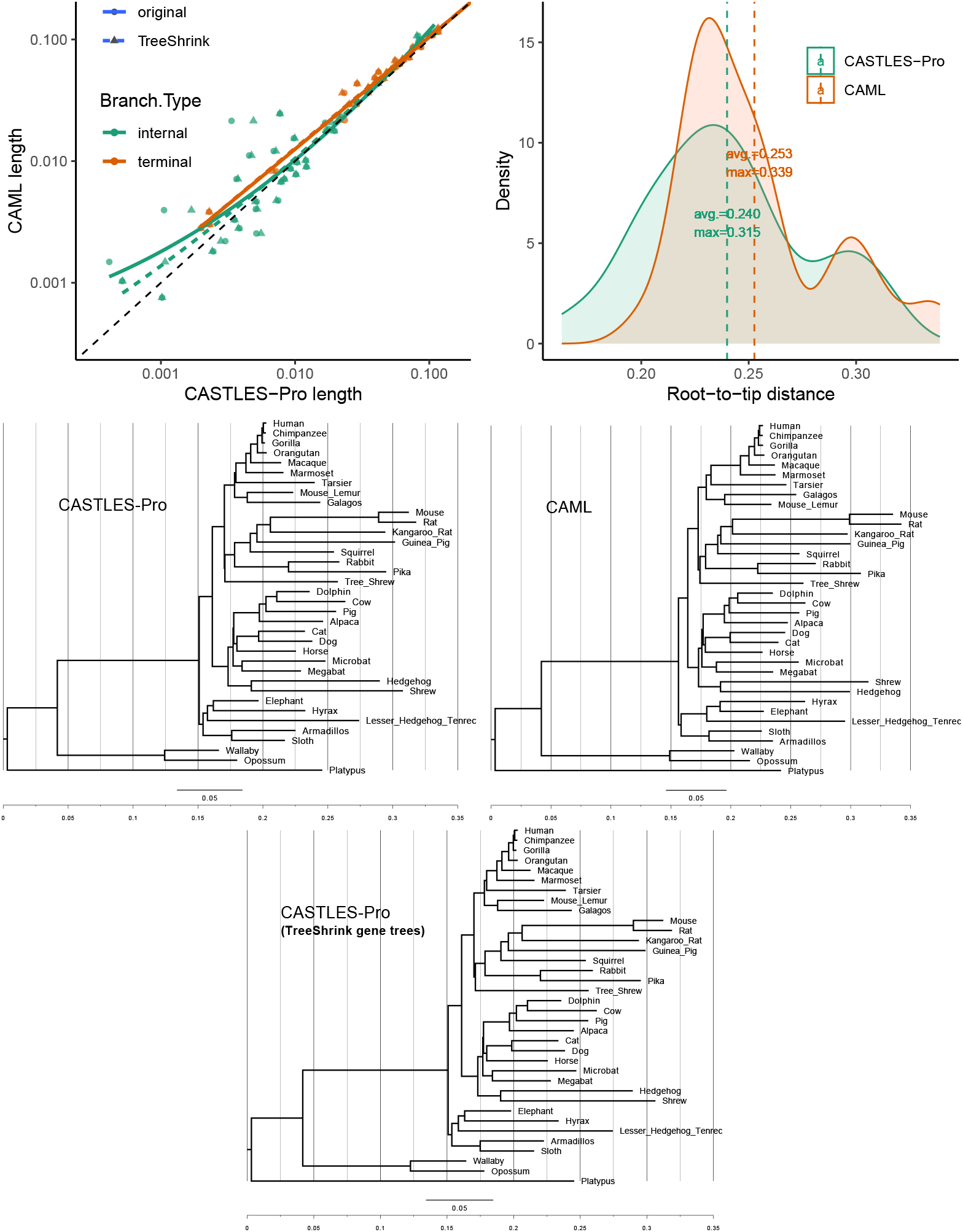
Comparison between branch lengths of CASTLES-Pro (with or without TreeShrink) and CAML on the mammals datasets of Song et al. (2012). The number of species is 37 and the number of gene trees is 424. The average gene sequence length is 3099bp. We draw branch lengths using CASTLES-Pro and concatenation on an ASTRAL species tree topology. The branch lengths are drawn after removing the outgroup taxa *Chicken*.

**Figure S18:**
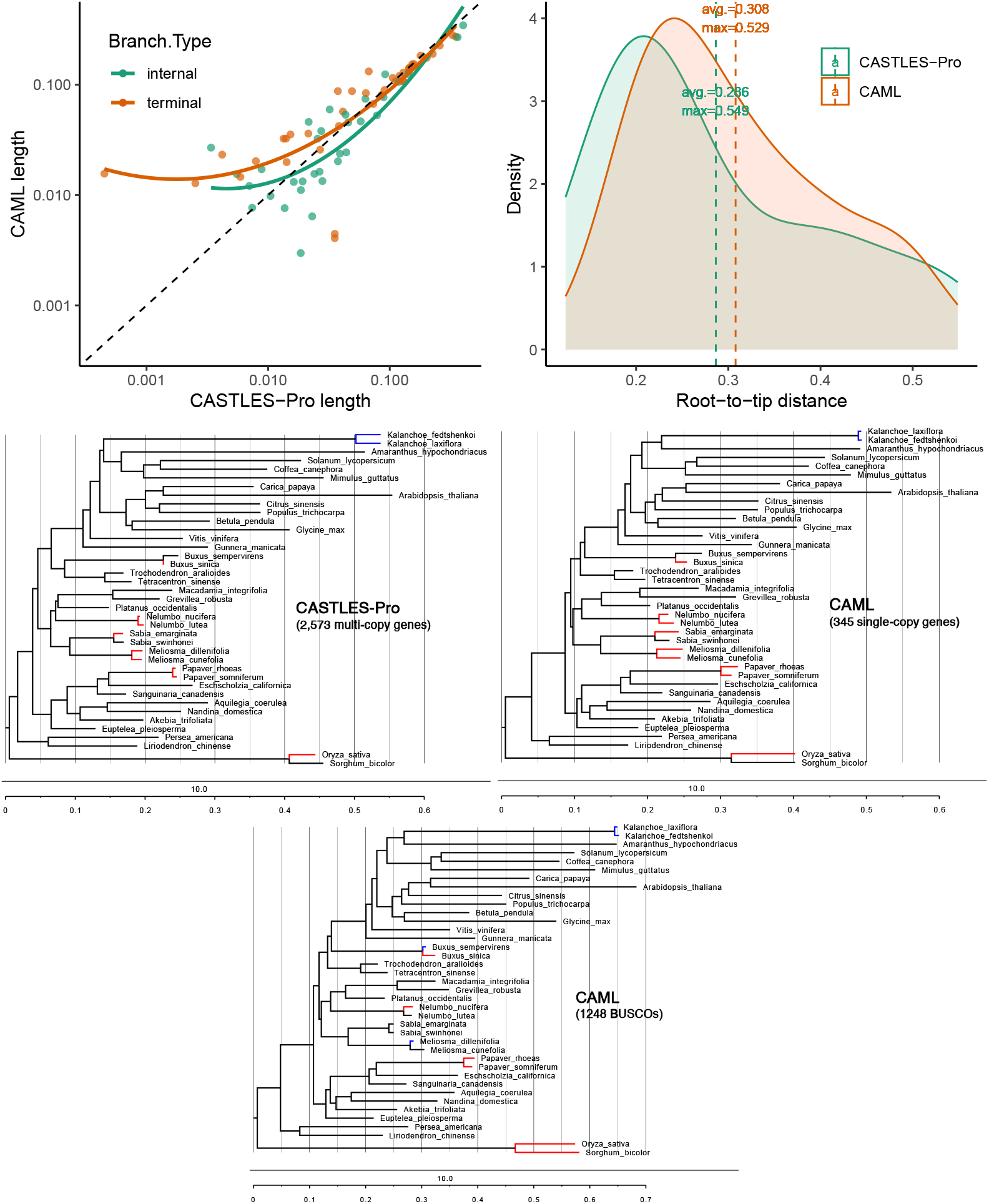
Comparison between branch lengths of CASTLES-Pro and CAML based on Angioseprm353 genes and BUSCO genes on the 40-taxon Eudicots datasets of Chanderbali et al. (2022) (after removing the outgroup *Amborella trichopoda*). The two CAML trees are estimated from the concatenation of 345 filtered Angioseprm353 single-copy genes or 1248 BUSCO single-copy genes, and CASTLES-Pro uses the 2,753 multi-copy gene family trees. The coalescent tree is estimated using ASTRAL-Pro2 and is different in 3 branches with the two concatenation trees (which are identical). The branch lengths that are at least 2x longer or shorter in CASTLES-Pro tree compared to the concatenation trees are highlighted in blue and red respectively.

**Figure S19:**
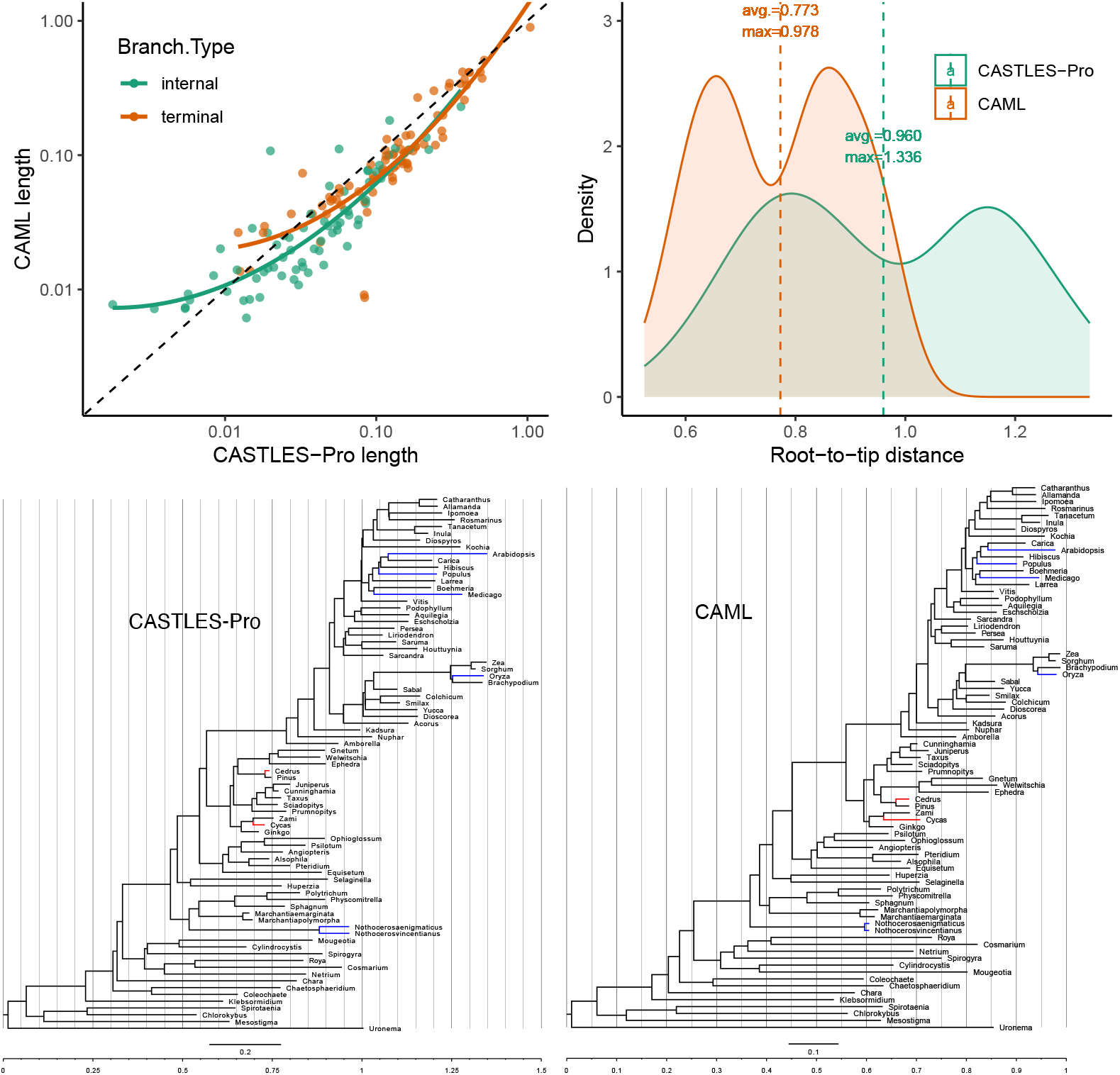
Comparison between the branch lengths of CASTLES-Pro and CAML on the plants dataset of Wickett et al. (2014). The CAML tree is from the original study and was estimated from the concatenated alignment of 424 single-copy genes. In addition, we estimated a tree from the set of 9,610 multi-copy gene family trees from that study using ASTRAL-Pro2 and used CASTLES-Pro to draw branch lengths on this tree. The taxa in the two sets of genes are not entirely identical (see Wickett et al. (2014) for more detail). The RF distance between the CAML and ASTRAL-Pro trees on 79 shared taxa is 9.2%. The correlation is reported only for the shared branches between two trees. The branch lengths that are at least 2x longer or shorter in CASTLES-Pro tree compared to the concatenation tree are highlighted in blue and red respectively.

**Figure S20:**
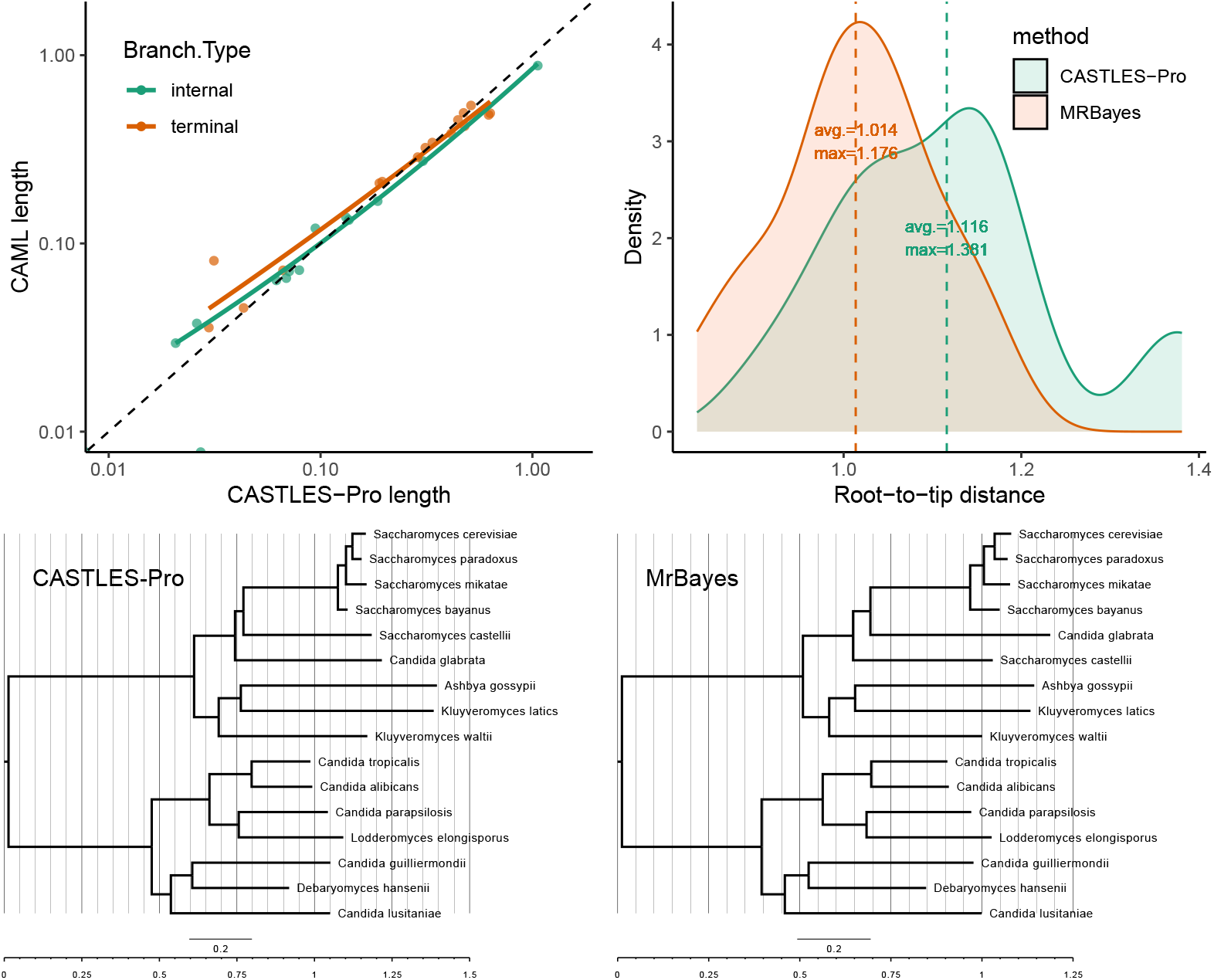
Comparison between branch lengths of CASTLES-Pro and concatenation with MrBayes on the fungal datasets of Butler et al. (2009). The number of species is 16. The original study had used MrBayes (Huelsenbeck and Ronquist, 2001) on a concatenated alignment created by sampling 30,000 sites from 706 individual gene family orthologous peptide sequences. We used ASTRAL-Pro2 to estimate a species tree using all 7,180 gene family trees, and used CASTLES-Pro to estimate branch lengths on that tree. The two trees are different in one branch, with an RF distance of 7.6%.

**Figure S21:**
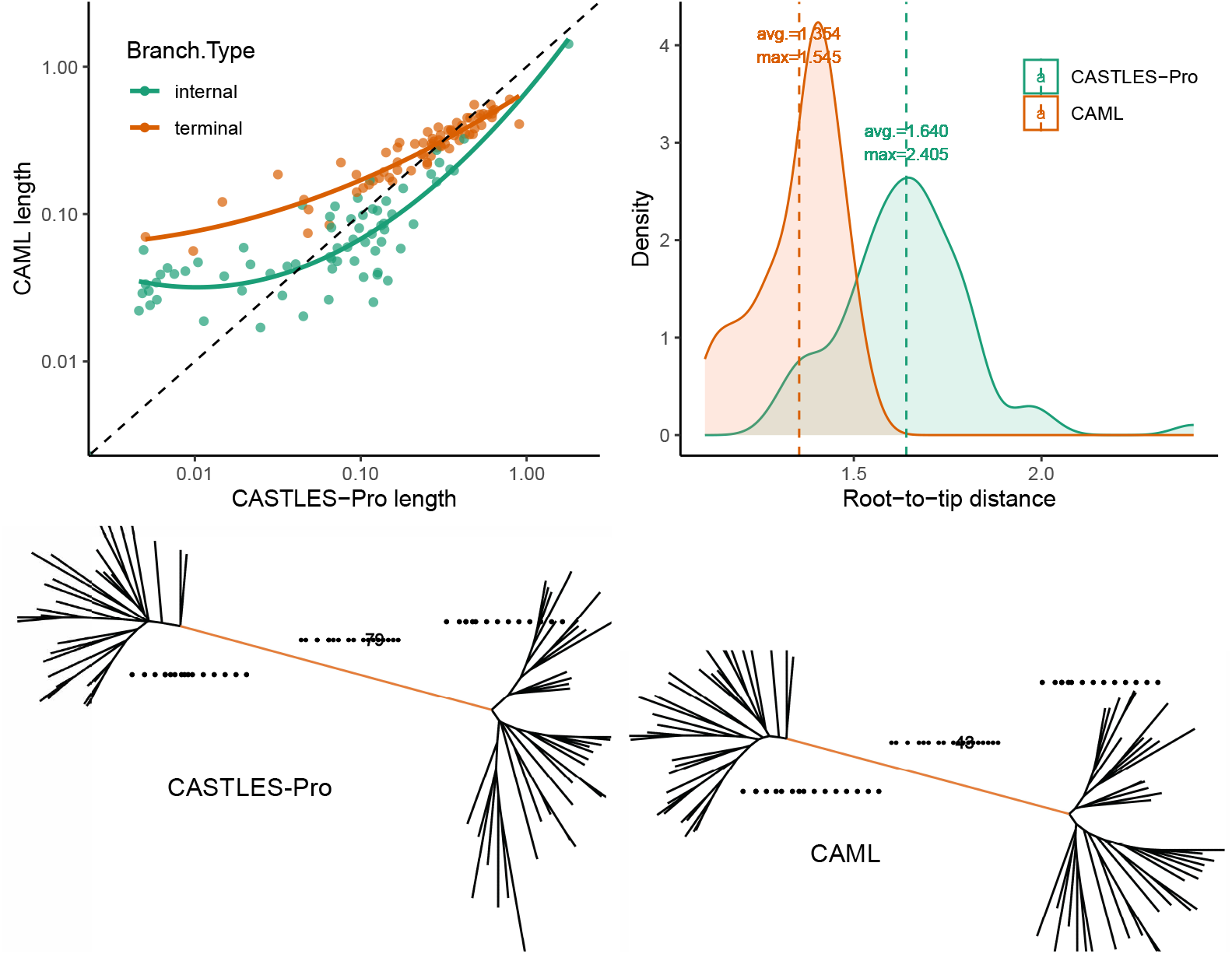
Comparison between branch lengths of CASTLES-Pro and CAML on the bacterial dataset with core genes from Williams et al. (2020). The number of species is 72 and the number of genes is 49. Both methods draw branch lengths on an ASTRAL tree topology. The branch colored in orange is the AB branch.

**Figure S22:**
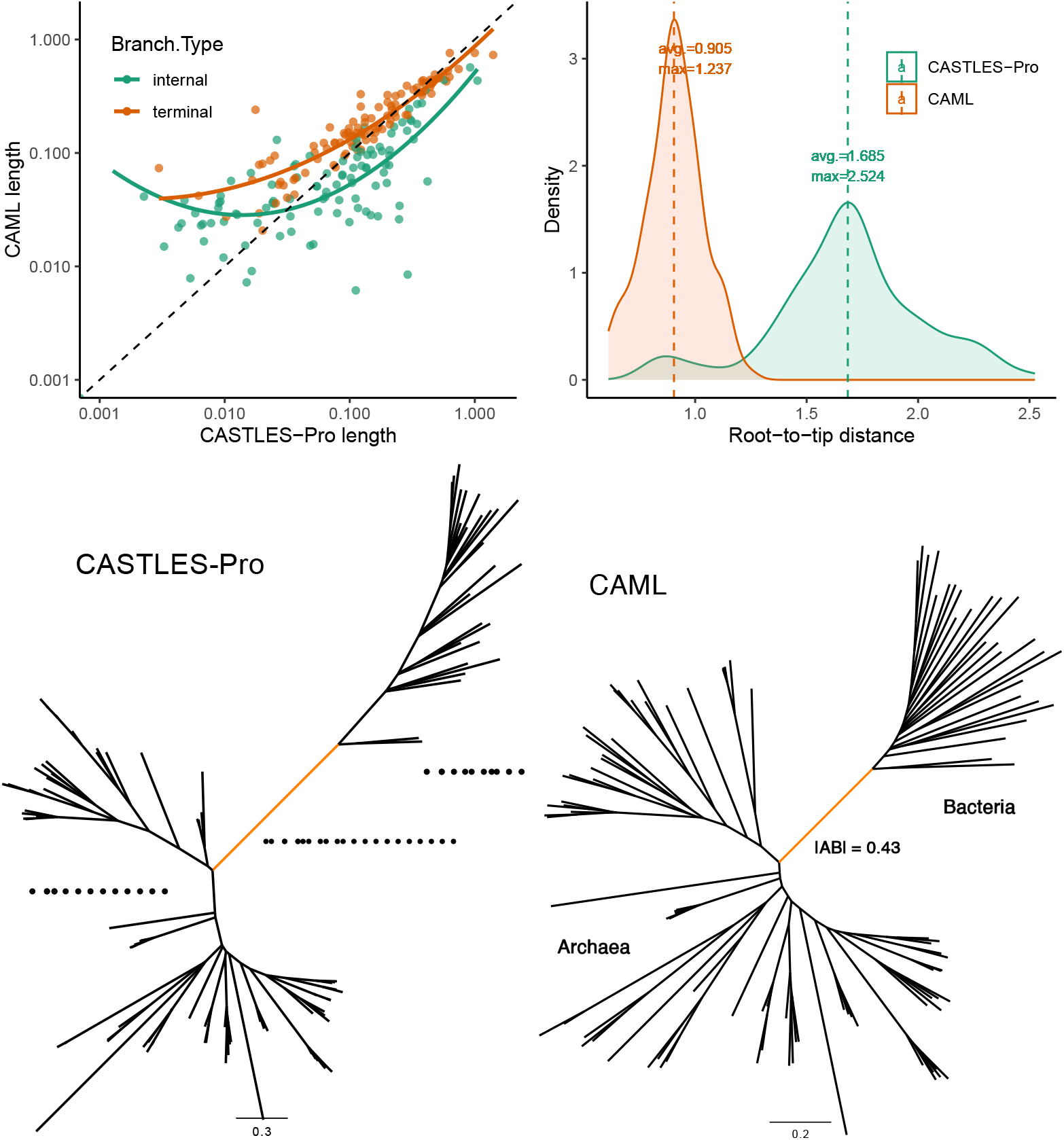
Comparison between branch lengths of CASTLES-Pro and CAML on the bacterial dataset with non-ribosomal genes from Petitjean et al. (2015). The number of species is 108 and the number of genes is 38. Both methods draw branch lengths on an ASTRAL tree topology. The total alignment length is 6,534bp. The branch colored in orange is the AB branch.

**Figure S23:**
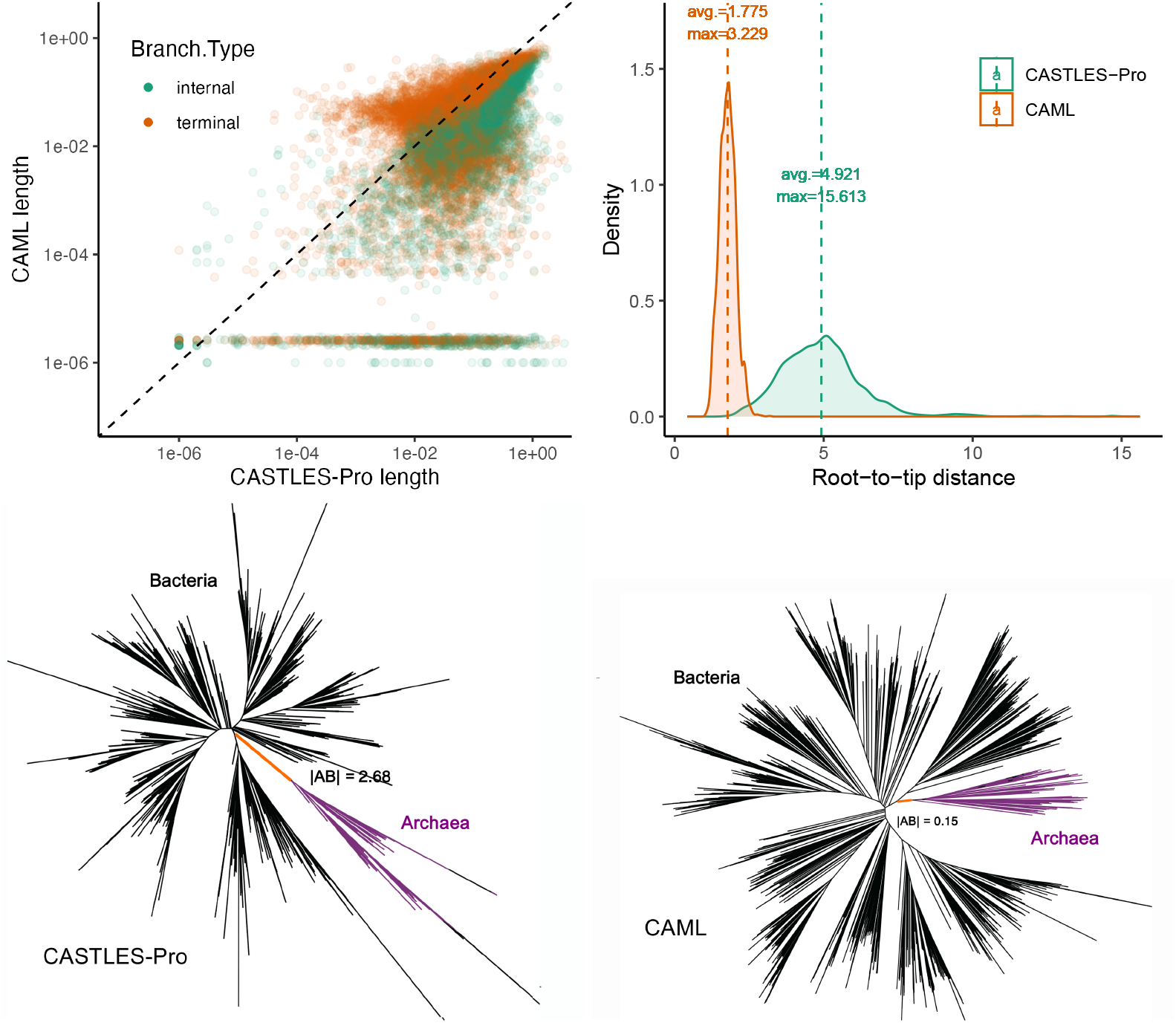
Comparison between branch lengths of CASTLES-Pro and CAML on the Web of Life (WoL) bacterial dataset from Zhu et al. (2019). The number of species is 10,575 and the number of marker genes is 381. The original study had estimated an ASTRAL tree topology and furnished that with concatenation branch lengths, using a concatenated alignment of size 38kbp. This alignment was created by selecting 100 random sites from each gene sequence, and was 5X shorter than the full-length alignment that had 192k sites in total. We draw branch lengths on the same tree topology using CASTLES-Pro. The long tail of branches with 1e-6 or 2e-6 length in the concatenation tree correspond to no-event branches. The branch colored in orange is the AB branch.

**Figure S24:**
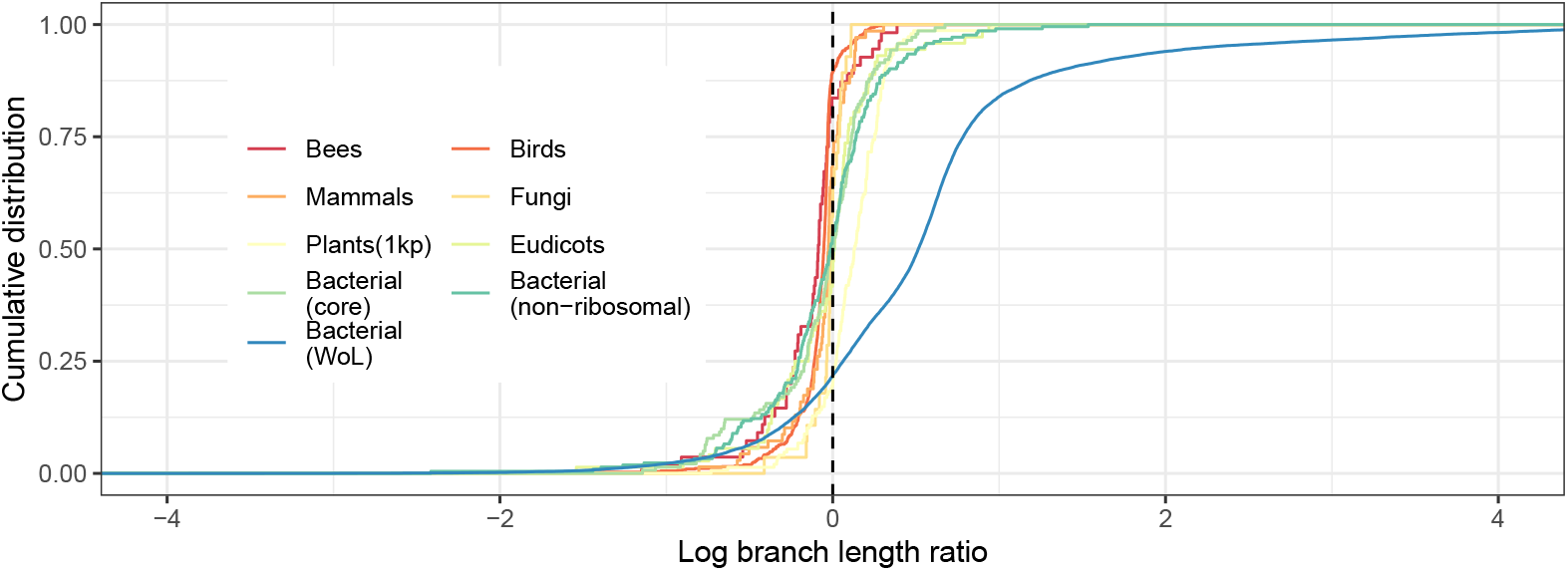
Cumulative distribution of the log ratio between the branch lengths produced by CASTLES-Pro and branch lengths produced by concatenation on nine biological datasets with different sources of gene tree heterogeneity.

**Table S10:** Runtime and peak memory usage of CASTLES-Pro on the biological datasets. Branch lengths are estimated on a fixed species tree topology. CASTLES-Pro uses multi-copy gene trees for the fungi, plants, and eudicots dataset and single-copy gene trees for the rest of the datasets.

| Dataset | # taxa | # genes | time (seconds) | peak memory (GB) |
| --- | --- | --- | --- | --- |
| Neoavian birds | 363 | 63,430 | 2034.64 | 54.83 |
| Bees (subfamily Nomiinae) | 32 | 853 | 1.85 | 0.06 |
| Mammals | 37 | 424 | 8.60 | 0.05 |
| Fungi | 16 | 7,180 | 1.60 | 0.20 |
| Plants (1kp) | 83 | 9,610 | 35.92 | 3.16 |
| Eudicots | 40 | 2,573 | 26.28 | 0.63 |
| Bacterial (core genes) | 72 | 49 | 1.72 | 0.01 |
| Bacterial (non-ribosomal genes) | 108 | 38 | 1.89 | 0.01 |
| Bacterial (WoL) | 10,575 | 381 | 3180.94 | 14.48 |

